# Insilico Investigation of Terpenoid efficacy on Cannabinoid Receptors using QSAR models and fragment-based Pharmacophore modelling

**DOI:** 10.1101/2025.09.04.674146

**Authors:** Aniruddhya Mukherjee, Khushhali Menaria Pandey

## Abstract

*Cannabis Sativa* a medical plant rich in phytochemicals have a range of terpenoids and cannabinoids with a wide range of therapeutic uses. The possible interaction of terpenoids with cannabinoid receptors implicated in analgesic pathways has not received enough attention. This work used in silico methods to target Cannabinoid Receptors Type I (CNR1) and II (CNR2) in order to find novel terpenoid compounds from Cannabis sativa that may have pain-relieving properties. QSAR modelling based on known cannabinoid receptor inhibitors was used to build and screen a curated library of 119 terpenoids. The terpenoid library was screened using pharmacophore models that were created. Molecular docking was performed on the top candidates using CNR1 (PDB: 5U09) and CNR2 (PDB: 5ZTY). To evaluate the stability of receptor–ligand complexes over 100 ns, molecular dynamics simulations were run. SwissADME was used for ADMET profiling in order to assess drug-likeness and pharmacokinetic characteristics. Both receptors’ QSAR models showed strong predictive power (CNR1: r2 = 0.854; CNR2: r2 = 0.798). γ-Eudesmol and Bisabolol were the top hits for CNR1 and CNR2, respectively, according to pharmacophore screening. Strong binding affinities were demonstrated by molecular docking (γ-Eudesmol: –7.9 kcal/mol; Bisabolol: –8.5 kcal/mol), and simulations verified the stable connections. Both compounds were drug-like, according to ADMET analysis, while γ-Eudesmol had better synthetic accessibility and fewer structural alarms. According to the results, bisabolol and γ-Eudesmol show promise as cannabinoid receptor-targeted analgesics. The finding encourages more biological validation and emphasises the potential of terpenoids obtained from Cannabis sativa in medicinal medication discovery.

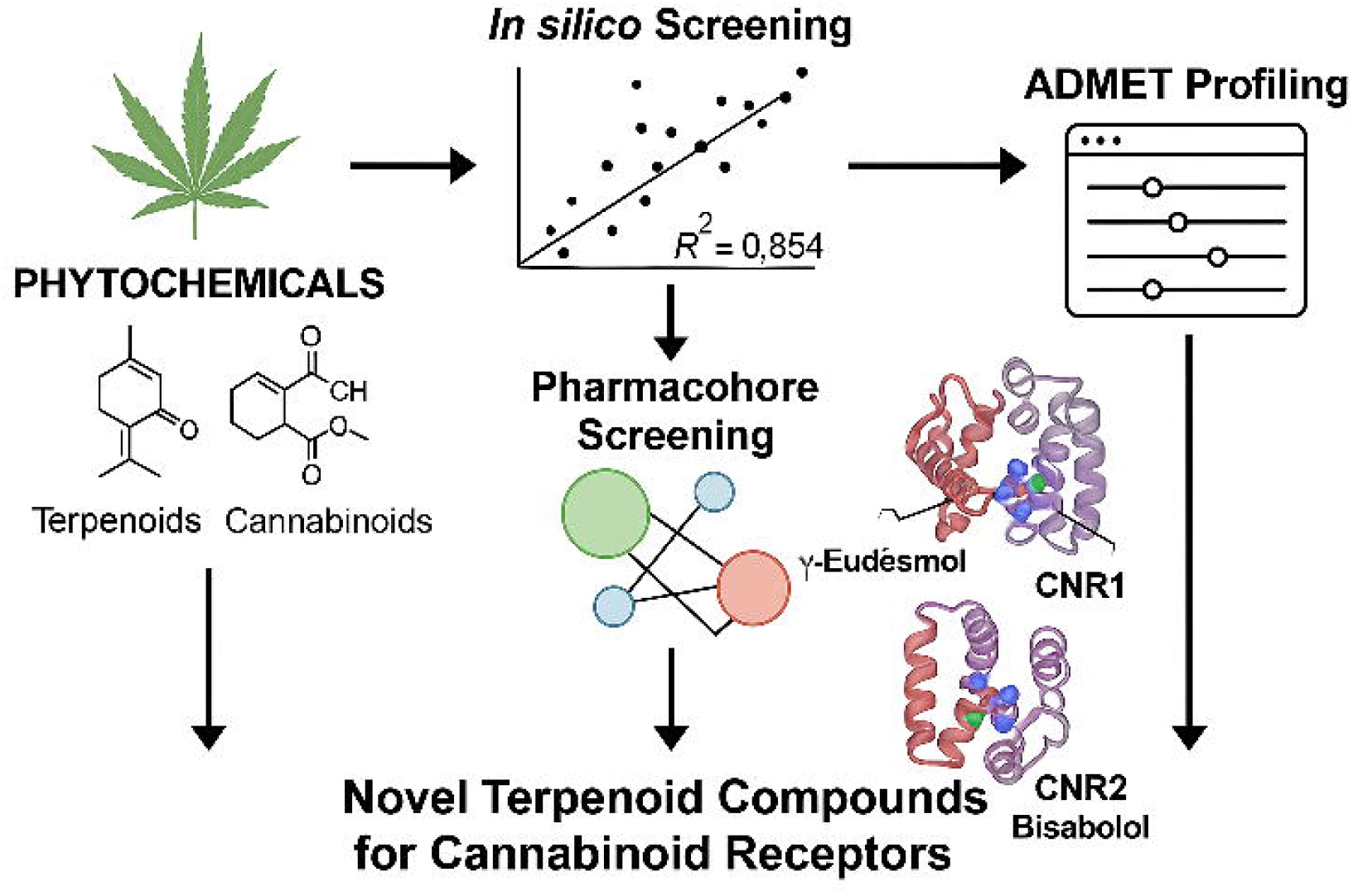

## INTRODUCTION

Since ancient times, Indian hemp, also known as Cannabis sativa L. (C. sativa), has been grown mostly in Central Asia (India and China)^[1]^. Over the years, Cannabis sativa has been utilised for recreational and religious purposes, as well as a source of fibre, food, oil, and medicine^[2]^. Cannabis sativa is a well-known medicinal plant that has attracted attention over the years and most recently because of the abundance of phytochemicals in Cannabis sativa (L.), which has long been used medicinally^[3]^, scientists have been working to unlock the plant’s pharmacological potential. Natural products extracted from Cannabis sativa (L.) can be categorized mainly into three classes: Major Cannabinoids like ▲9-THC (Tetrahydrocannabinol)^[4]^, ▲8-THC^[5]^, CBG (Cannabigerol)^[4]^, CBC (Cannabichromene)^[6]^, CBD (Cannabidiol)^[7]^, CBND (Cannabinodiol)^[8]^, CBE (Cannabielsoin)^[9]^, etc. are phytochemicals that are predominantly found (structures shown in Supplementary Fig 1), Minor Cannabinoids like ▲9-THCV (Tetrahydrocannabivarin)^[10]^, CBGA (Cannabigerolic Acid)^[11]^, CBLV (Cannabicyclovarin)^[12]^, etc are chemical analogues of major cannabinoids extracted from different strains of Cannabis sativa on basis of geographical diversities (structures shown in Supplementary Fig 2). Terpenoids or isoprenoids, comprise the second-largest class of cannabis chemical components. These substances are in charge of giving the plant its distinctive smell. From Cannabis sativa, terpenoids such as (−) pinene, (−) terpinene, camphene, and linalool have been isolated^[13]^ (structures shown in Fig 1).

**Fig 1:**
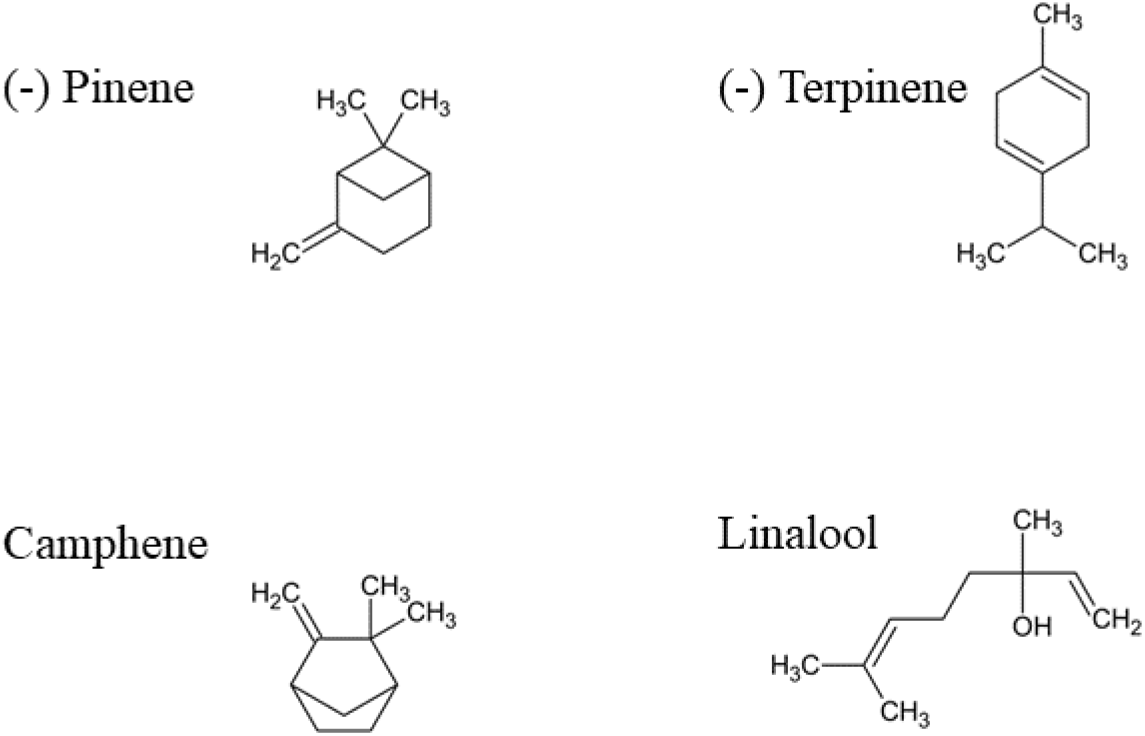
Terpenoids obtained from Cannabis sativa

As of 2017, chemical constituents of Cannabis sativa can be summarized as 125 cannabinoids, 42 phenolics, 34 flavonoids, 120 terpenes and 2 alkaloids. One of the G-protein-coupled receptors with the greatest expression in the brain is known to be the cannabinoid receptor type 1 (CNR1) receptor. One way they work is through an extremely common kind of retrograde signalling called endocannabinoid-mediated depolarization-induced suppression of inhibition, in which depolarization of a single neuron results in a decrease in GABA-mediated neurotransmission. Cannabinoid receptor type 2 (CNR2) is found on immune system T cells, macrophages, and B cells, as well as in haematological cells, the brain, and the central nervous system (CNS). In keratinocytes, they serve a purpose as well. Peripheral nerve terminals also express them. Antinociception, or the alleviation of pain, is facilitated by these receptors. Their function in the brain, where they are mostly expressed by microglial cells, is still unknown. The primary targets for cannabinoids, terpenoids and endocannabinoids are these cannabinoid receptors. Cannabinoids have a diverse medical therapeutic potential ranging from antibacterial, antifungal, and anticancer properties. Murine collagen-induced arthritis (CIA) was used to test the therapeutic potential of cannabidiol (CBD), the main non-psychoactive component of cannabis by^[14]^. Antibacterial cannabinoids found in Cannabis sativa have long been recognized to exist, but their potential to combat antibiotic resistance has not previously been explored. Appendino and his team in 2008 experimented on five main cannabinoids (cannabidiol, cannabichromene, cannabigerol, 9-tetrahydrocannabinol, and cannabinol)^[15]^ all showing strong effectiveness against a range of MRSA (methicillin-resistant Staphylococcus aureus) strains that are now relevant to clinical practice. Being a compound with such high biological activity and medicinal values cannabinoids have their own curse. The effects of THC on blood pressure and heart rate are immediate and dose-dependent. Higher doses and rising usage frequency are frequently seen as a result of a THC tolerance that develops quickly. There is evidence to show that frequent marijuana usage raises the risk of myocardial infarction (MI) and cardiac arrhythmias^[16]^. Mendizabal & Adler-Graschinsky in 2007 reported that long-term marijuana usage may cause ANS dysfunction^[17]^, which could result in cycles of vasoconstriction and vasodilation independent of skeletal muscle action. However, rigorous evidence-based clinical investigations and proven scientific mechanisms are lacking for topical borneol’s therapeutic usefulness. In a randomised, double-blind, placebo-controlled clinical investigation comprising 122 patients with postoperative pain, Wang et al^[18]^ evaluated the analgesic effectiveness of topical borneol. Their research shows that borneol, which the US FDA presently only approves for use as a food adjuvant or flavouring agent, is an efficient topical painkiller in humans and identifies a crucial component of the molecular mechanism behind its analgesic effect. Inflammation is one of the body’s defence mechanisms and is characterised by redness, swelling, pain, and a feeling of heat. Although the inflammatory response is crucial for the survival of the host, it can also result in chronic inflammatory disorders. Linalool is a naturally occurring chemical found in the essential oils of various types of aromatic plants. Along with other bioactive qualities, it has anti-inflammatory and antinociceptive effects. In the current work, using lipopolysaccharide (LPS)-stimulated RAW 264.7 cells and an in vivo lung damage model, Huo et al looked at the anti-inflammatory properties of linalool. They examined how linalool affected Raw 264’s production of inflammatory mediators after exposure to LPS^[19]^.

Even having such high standards of medicinal values medicinal cannabis has very little reach of research and very limited experimental data can be obtained. A lot more research needs to be carried out to find the possible potential of medicinal cannabis, biological pathways it is linked to and plausible therapeutic mechanisms for experimented medical activity. The current objective of this study is to identify novel terpenoids obtained from Cannabis sativa that can have possible pain killing activity demonstrated using computational approaches.

## Methodology

### Protein selection and Structure preparation

Most of the insilico studies that have been carried out on cannabinoid receptors have been carried out for High-resolution crystal structure of the human CNR1 cannabinoid receptor (PDB: 5U09)^[20]^ and Crystal structure of human G protein-coupled receptor (PDB: 5ZTY)^[21]^. So, both structures were downloaded from the Protein Data Bank (https://www.rcsb.org/). R-Value free was 0.238 and R-Value work was 0.180 with a resolution of 2.60L was selected for the type I receptor and R-Value free of 0.275 and R-Value work 0.224 with a resolution of 2.80□ was selected for type II cannabinoid receptor. To prepare the macromolecules and other co-existing water molecules and non-standard residue were removed using Biovia Discovery Studio 2021 Client. To produce a protonated state at physiological pH, one uses Autodock4.2, build-up geometry optimization, and the addition of polar hydrogen Kollman charges and Gasteiger charges.

### Cannabinoid and Terpenoid Library preparation

Through a rigorous search of the literature, a total of 119 terpene compounds were obtained out of which 60 were monoterpenes, 51 were sesquiterpenes, 2 diterpenes, 2 triterpenes, and 4 miscellaneous terpenes (Supplementary Table 1). All the compounds were downloaded from PubChem (https://pubchem.ncbi.nlm.nih.gov/). Another set of compounds with experimental activity was downloaded for QSAR and Pharmacophore analysis. From PubChem Bioassay cannabinoids with verified biological activity against Cannabinoid Receptor Type I (62 compounds) (Supplementary Table 2) and II (67 compounds) (Supplementary Table 3) were downloaded.

### Creation of Partial Least Square QSAR models

Individual QSAR models were generated for compounds having verified biological activity for Cannabinoid Receptor Type I and Type II using the Partial Least Square algorithm in the Accelrys Discovery Studio 3.1. Individual sets of compounds downloaded from the database were loaded into the system and compiled into a single file along with their biological activity (Ki or inhibitory constant in this case). The inhibitory constant for all the compounds had a diverse range of valuation which would hamper the models so normalization needed to be performed on the data. The biological activity values were log-transformed using MS Excel and added as an attribute in Discovery. Log transforming the values normalized the diverse ranging values into a definite range for model prediction. Then the compounds were prepared (i.e., standardizing charges for common groups, retaining the largest fragment, adding Hydrogen, representing ligands in Kekule from, generating a 3D geometry, removing duplicates, enumerating ionization states, and generation of canonical tautomer) for model prediction using the “Prepare Ligands for QSAR” tool. Then the prepared compounds were split into training and test set using the tool “Generate Training and Test Data”. In the course of QSAR model prediction compounds have to be divided into two groups, one is the Training set which is used for model prediction and the other one is the Test set which is used to validate the quality of the QSAR model predicted. After the generation of the training and test set using the “Create Partial Least Square Model” tool, the QSAR model was predicted and cross-validated using the test set as well.

### Pharmacophore Model Generation and Ligand Screening

The best QSAR model was chosen and pharmacophore features like HBA, HBD, Hydrophobic, Ring Aromaticity, etc were investigated using the “Feature Mapping” tool. Using those features a suitable pharmacophore was built using the “Auto Pharmacophore Generation” tool. A total of 10 pharmacophore models were set to be produced and the pharmacophore with the greatest number of features was to be chosen for further analysis.

The best feature-mapped pharmacophore was chosen and screened against our designed terpenoid library using the tool “Screen Library” which is used to map multiple ligands to a single pharmacophore model. The top three compounds with the highest fit values were considered for further Molecular Docking Analysis.

### Molecular Docking

Molecular Docking was performed using AutoDock Vina^[22]^. Upon review of the literature specific site of activity or ligand interaction for both the Cannabinoid receptors were known. So, the screened terpenoids were converted into PDBQT using the OpenBabel program^[23]^ and molecularly docked with the cannabinoid receptors at their respective active sites. The results of docking that is the protein-ligand complex interaction were visualized using Discovery Studio Visualizer 2021.

### Molecular Dynamic Simulation Studies

The best-performing terpenoid on the basis of binding affinity and binding mode was simulated with its respective receptor. For simulation, GROMACS 2021^[24]^ was used and for ligand topology generation SWISS Param (https://www.swissparam.ch/)^[25]^ was utilized. Forcefield was set to be CHARMM27^[26]^ for both the protein and protein-ligand complex. For protein, the water model used was SPC216 and for protein-ligand complex water model chosen was TIP3. The salt type for both the apoprotein and complex molecule was set to be NaCl. Energy minimization was set according to the Verlet cut-off scheme with a maximum of 50,000 minimization steps. For volume and pressure, the equilibration time step for integration was 2fs and the system was equilibrated for a total of 50,000 steps. For the final step, the time step was the same but the system for both the apoprotein as well as the complex was simulated for 50,000,000 steps resulting in a total simulation time of 100ns.

### ADMET Toxicology Analysis

A detailed tabulation of the ADMET properties including the pharmacokinetic properties and toxicological characteristics of the terpenoids was obtained from SWISS ADME (http://www.swissadme.ch/)^[27]^.

## Results and Discussion

### Creation of Partial Least Square QSAR Models

Active compounds against CNR1 and CNR2, 61 compounds and 66 compounds were retained. The CNR1 set was divided into 43 compounds in the training set and 18 compounds in the test set and subsequently the CNR2 set was divided into 46 compounds being in the training set and 20 compounds in the test set s shown in Fig 2 and 3 respectively.

**Fig 2:**
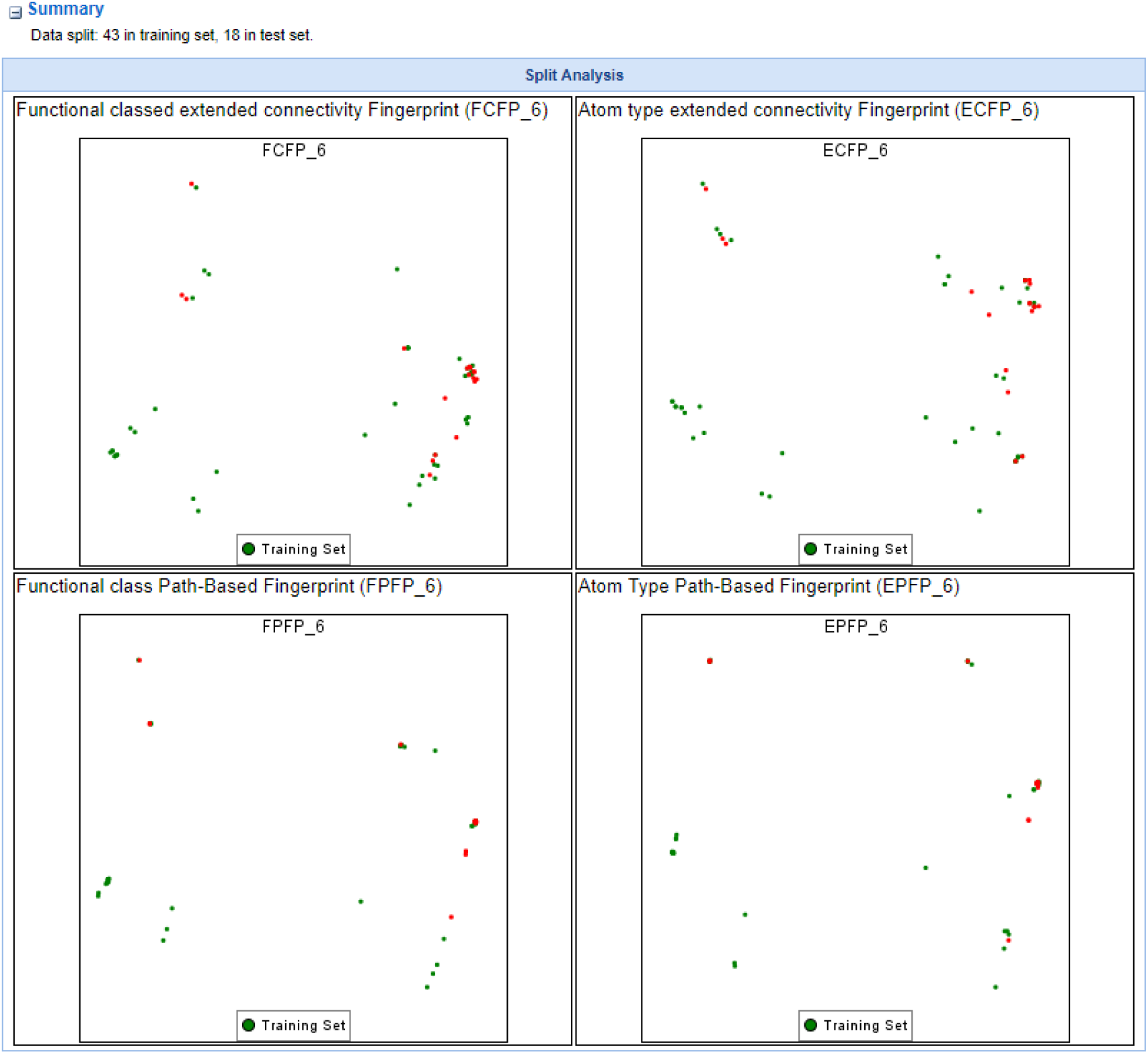
Training set generation for CNR1 active compounds

**Fig 3:**
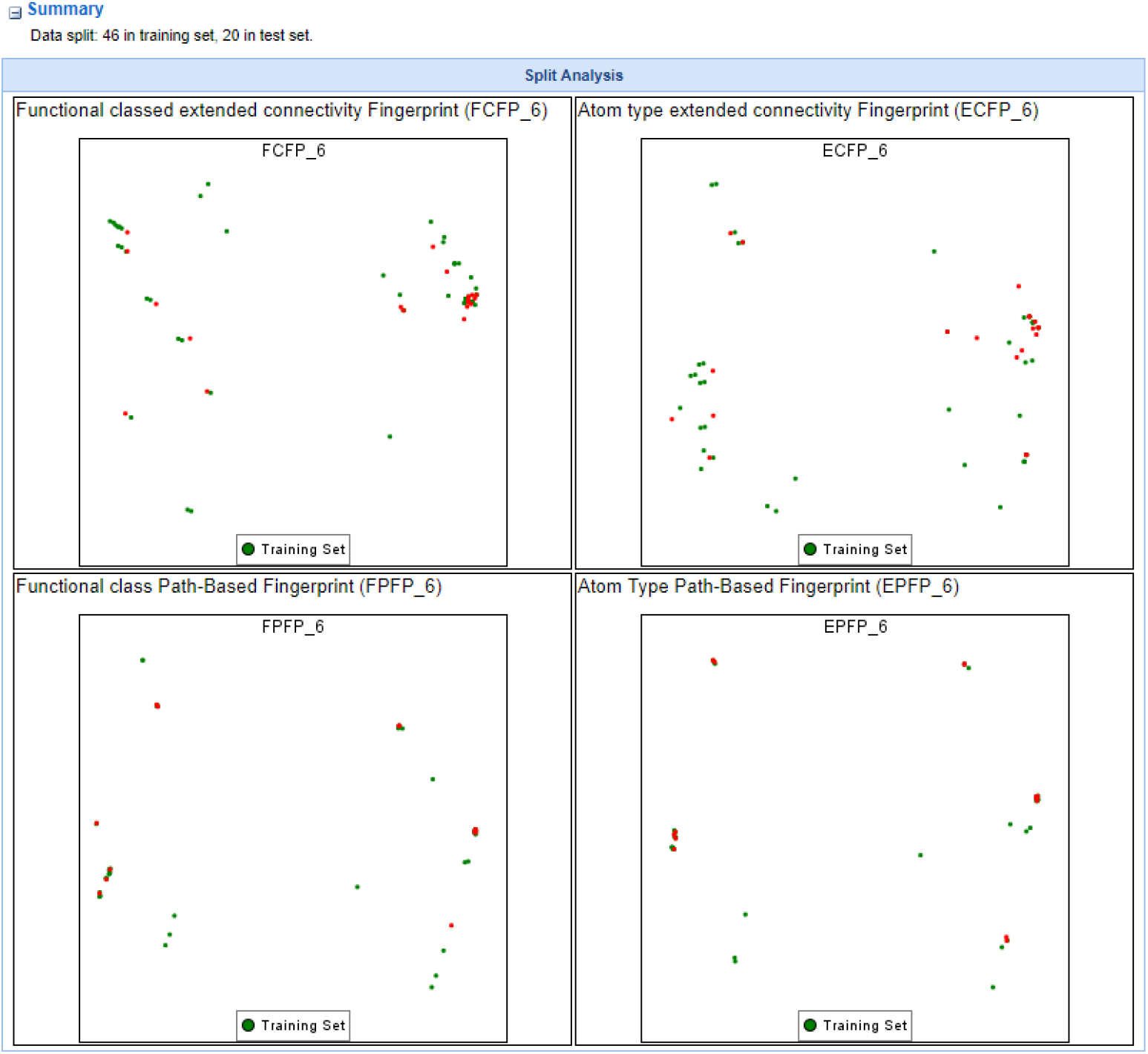
Training set generation for CNR2 active compounds

The results from the training set of both the receptors indicate that the final model prediction is going to be fairly well because the more clustered the data is in the training set generation better the predicted model is. As we can see in Fig 2 and 3, data in both the figures are fairly clustered, so we cannot expect a very good QSAR model. The QSAR model predicted for CNR1 has a r2 value of 0.854 as shown in Fig 4 which is actually a good score, r2 values ranging from 0.7 to 0.9 is considered as to be a sign for a good predicted QSAR model.

**Fig 4:**
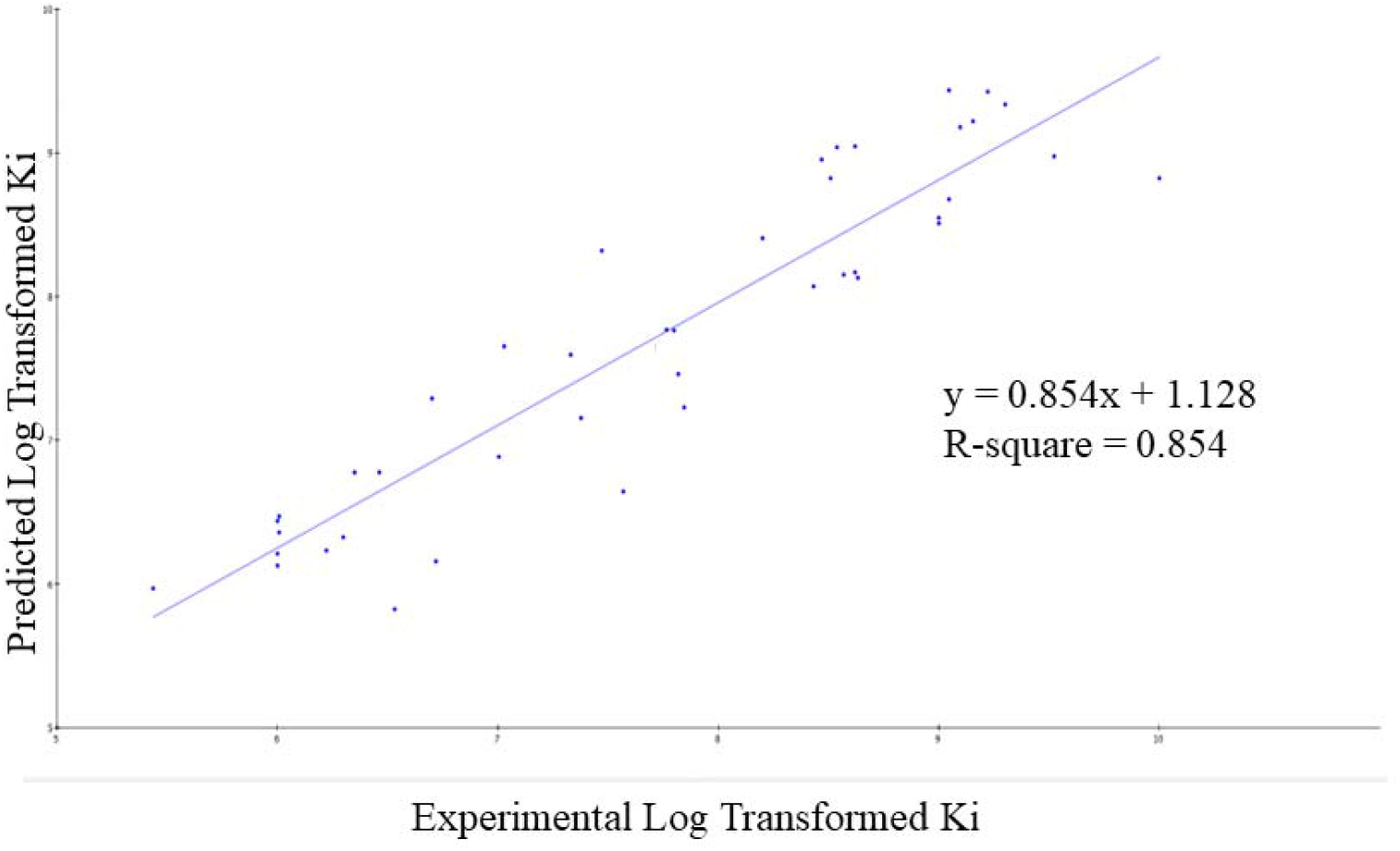
2D-QSAR model predicted for active compounds against CNR1

However, in case for CNR2 the r^2^ value came out to be 0.798 as shown in Fig 5 a bit low in the 0.7 to 0.9 range of model validation but a fairly good model. The predicted values of activity from the QSAR modelling for CNR1 and CNR2 are shown in table x and y respectively (Supplementary Table 4, 5). From the graph and the tables above, we get to know that PubChem CID 66941822 for CNR1 and PubChem CID 89994947 for CNR2 are two compounds which is closest to the model respectively, thus these two compounds were considered for pharmacophore generation.

**Fig 5:**
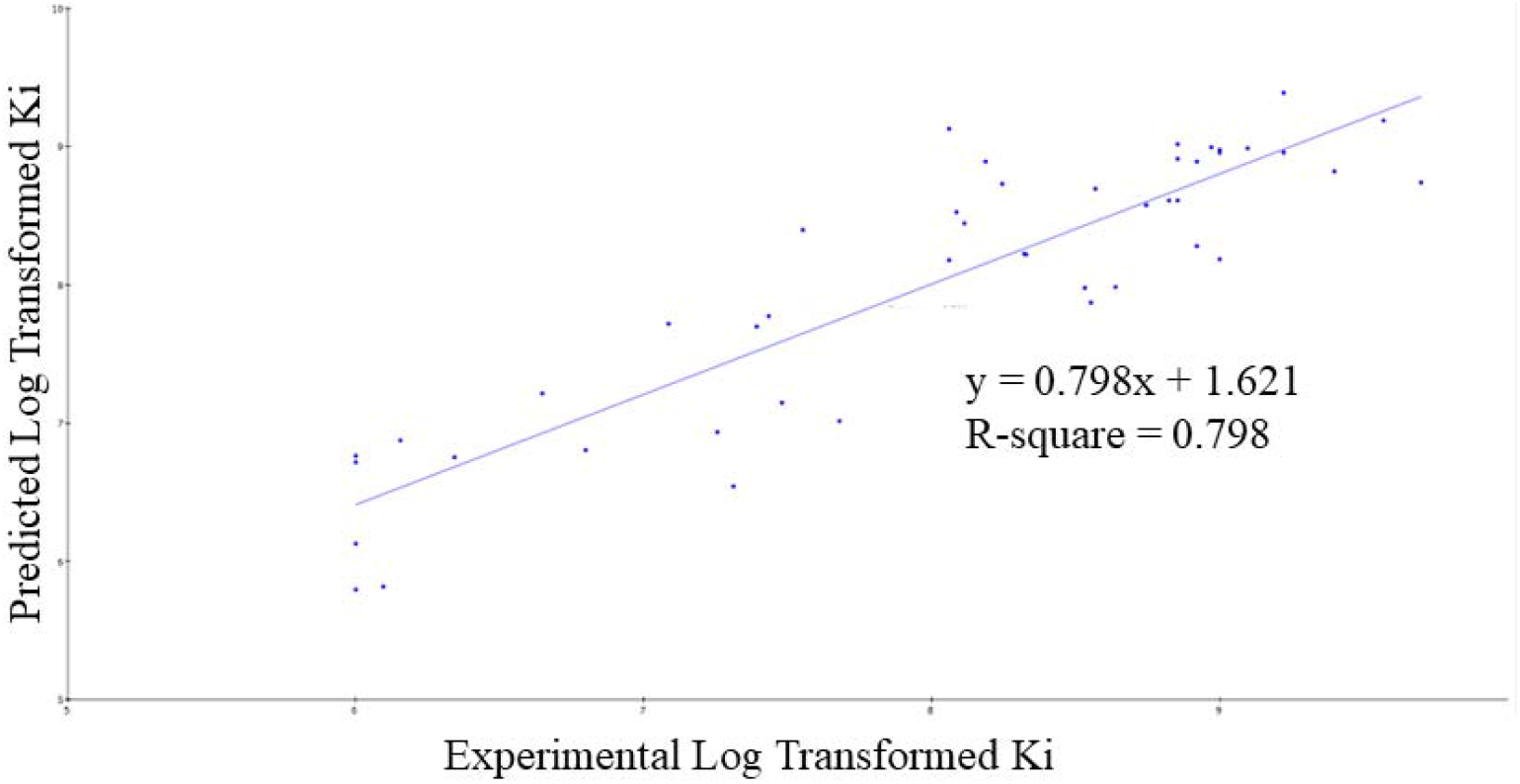
2D-QSAR model predicted for active compounds against CNR2

### Pharmacophore Generation and Library Screening

Pharmacophore using features were generated from CID 66941822 and CID 89994947 respectively as shown in Fig 6 and 7 respectively.

**Fig 6:**
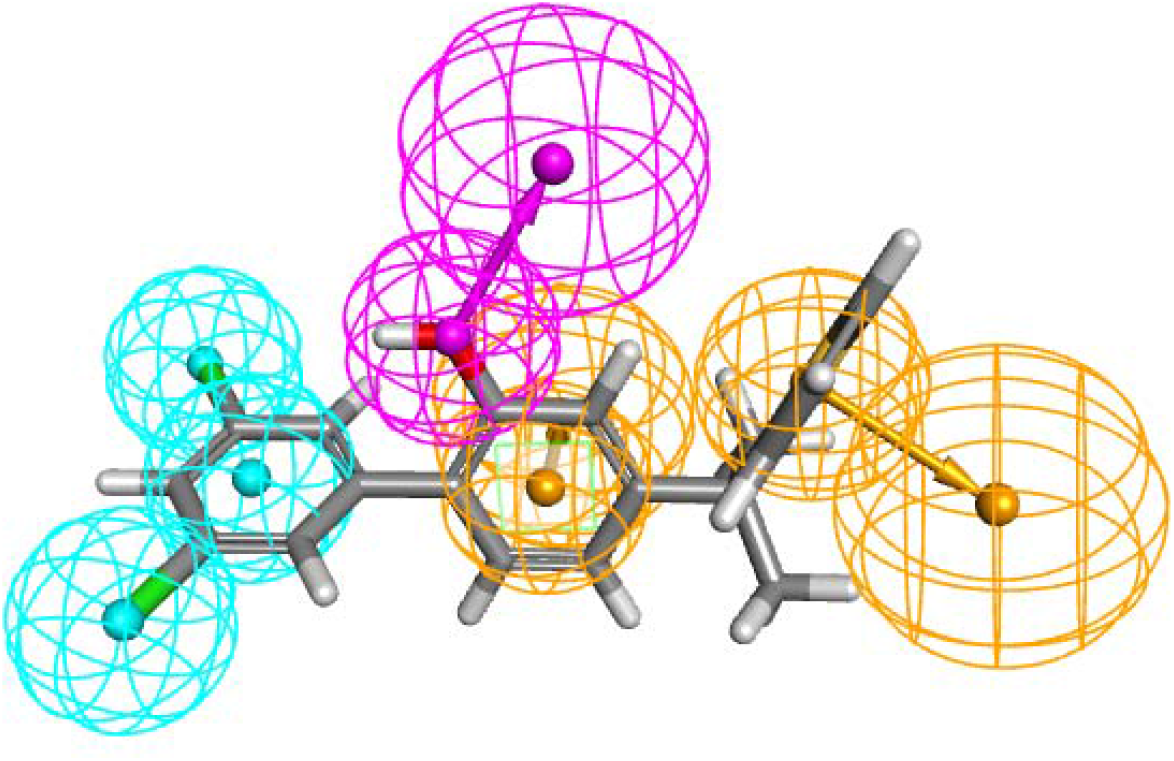
Pharmacophore model of CID 6694182 (Where Acceptor Feature is Green, Hydrophobic Feature is Cyan, Ring Aromatic Features are Orange, Doner Feature is Purple)

**Fig 7:**
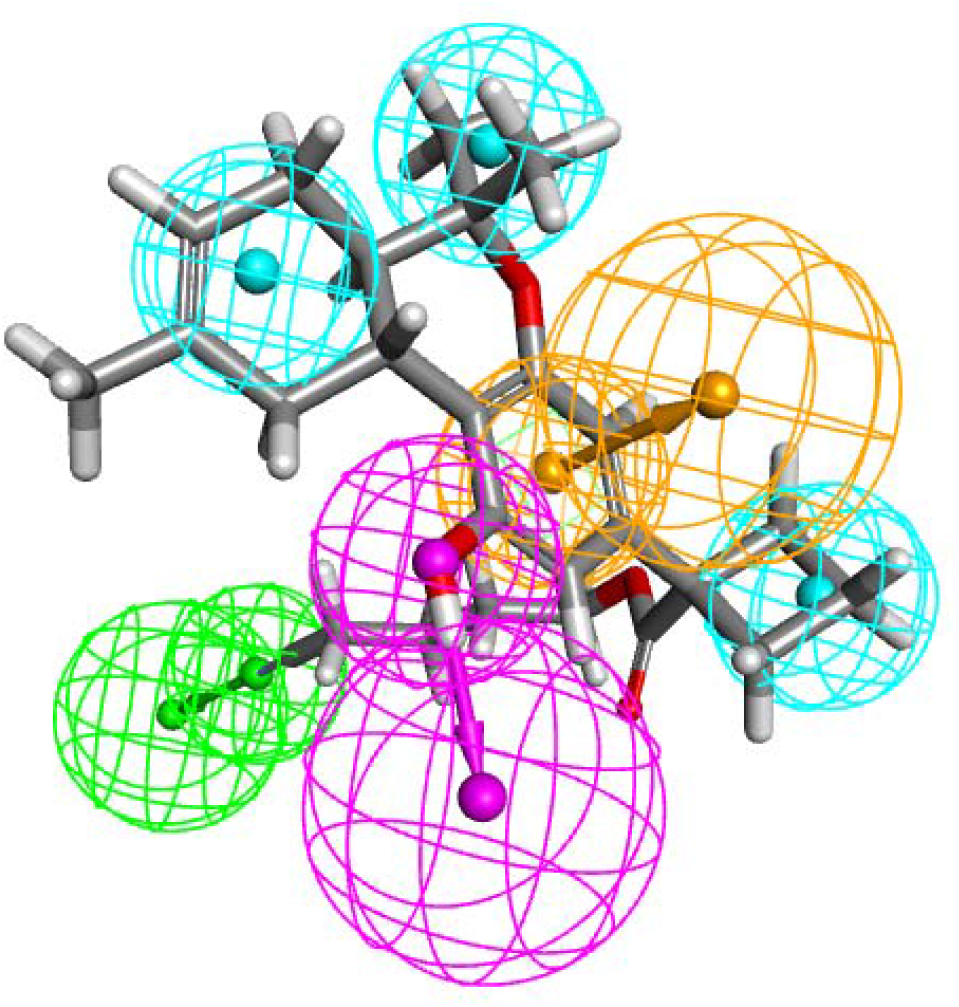
Pharmacophore model of CID 89994947 (Where Hydrophobic Feature is Cyan, Ring Aromatic Features are Orange, Where Acceptor Feature is Green, Doner Feature is Purple)

Upon screening these pharmacophores upon the self-designed terpenoid library, CID 66941822 was mapped upon 19 terpenoids (as shown in Supplementary Table VI, VII) and CID 89994947 was mapped on 11 terpenoids as shown in table. The molecule with the highest fit value was considered as the best mapping on the pharmacophore. From the table we can say that in case for CNR1 the top 5 mapped ligands are Guaiol (CID 227829), Nerol (CID 643820), Clovandiol (CID 76319362), (E, E)-Farnesol (CID 445070), Citronellol (CID 8842) and γ-Eudesmol (CID 6432005). And for CNR2 the top 5 mapped ligands are Nerolidol (CID 5284507), Carvacrol (CID 10364), α-Curcumene (CID 92139), Calamenene (CID 6429077), Bisabolol (CID 1549992).

### Molecular Docking

The top 3 terpenoids were used to perform Molecular docking for their respective cannabinoid receptors. Docking was performed on AutoDock Vina and the receptor-ligand interactions were viewed on Discovery Studio Visualizer 2021. Docking was performed on literature based active site residues for both the receptors individually and exhaustiveness of the experiment was set to be 10. The detailed information about binding affinity and binding mode are tabulated below in Table I and II for CNR I and CNR II respectively.

**Table 1:**
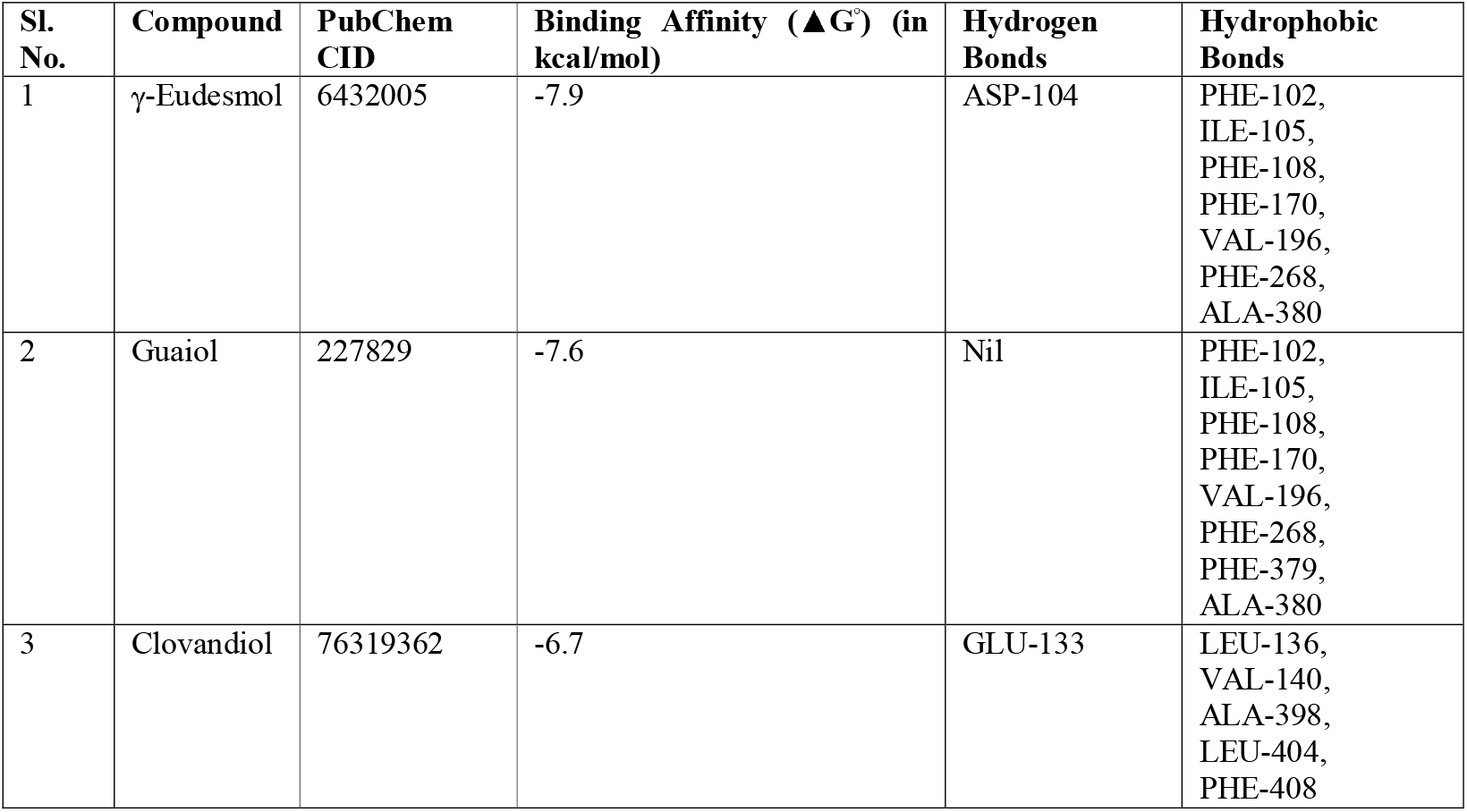
Binding affinities on CNR I along with Hydrogen bonding and Hydrophobic interactions.

**Table 2:** Binding affinities on CNR II along with Hydrogen bonding and Hydrophobic interactions.

| Sl. No. | Compound | PubChem CID | Binding Affinity ($\Delta G^\circ$) (in kcal/mol) | Hydrogen Bonds | Hydrophobic Bonds |
| --- | --- | --- | --- | --- | --- |
| 1 | Bisabolol | 1549992 | -8.5 | Nil | 110-ILE, 113-VAL, 117-PHE, 183-PHE, 186-ILE, 190-TYR, 191-LEU, 194-TRP, 258-TRP, 262-LEU |
| 2 | Calamene | 6429077 | -8.3 | Nil | 25-TYR, 87-PHE, 91-PHE, 94-PHE, 95-HIS (π-Stacking), 106-PHE, 110-ILE |
| 3 | α-Curcumene | 92139 | -7.5 | Nil | 87-PHE (π-Stacking), 91-PHE, 94-PHE, 106-PHE, 110-ILE, 113-VAL, 183-PHE, 281-PHE |

For CNRI we can see (Fig 8) that except for Clovandiol (−6.7 kcal/mol) both γ-Eudesmol (−7.9 kcal/mol) and Guaiol (−7.6 kcal/mol) have interacted with almost all the critical residues from the active site as shown in Fig 8. Residues like PHE-102, ILE-105, PHE-108, PHE-170, VAL-196, PHE-268, ALA-380 are common for both the terpenoids. In excess Gamma-Eudesmol has a Hydrogen bond interaction at an active site residue ASP-104 and Guaiol has an extra Hydrophobic interaction at PHE-379. The binding mode of Clovandiol is completely different than that of the other two terpenoids and do not match with any of the critical residues from the active site maybe due to its conformational rigidity.

**Fig 8:**
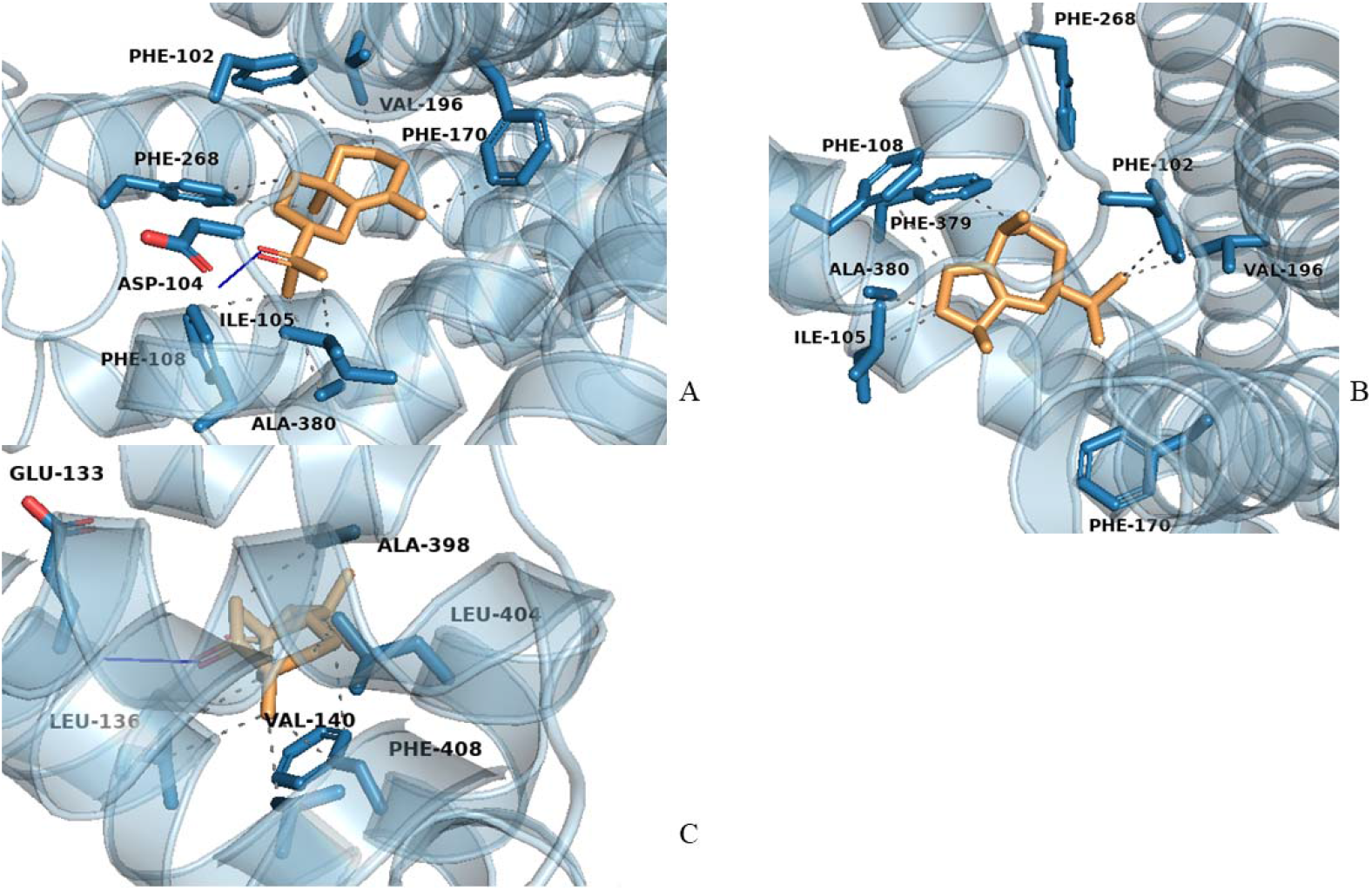
Protein ligand complexes formed by γ-Eudesmol (A), Guaiol (B) and Clovandiol (C) with CNRI (PDB ID: 5U09)

In case of CNRII we can see (Fig 9) that binding scores are elevated than CNRI, with Bisabolol scoring −8.5 kcal/mol, following Calamene scoring −8.3 kcal/mol and lastly α-Curcumene scoring −7.5 kcal/mol. It was expected that Bisabolol may have Hydrogen bonding like other ligands but when the complex was seen closely seen it was noticed that due to steric congestion and C-H bond conformations a stable Hydrogen bond was not established with the OH-group. And the rest of the compounds in the group are compounds devoid of heteroatoms thus incapable of Hydrogen-bond interaction. In case of binding mode and active site coverage all three compounds have hydrophobic interactions with almost all critical residues like 110-ILE, 113-VAL, 117-PHE, 183-PHE, 186-ILE, 190-TYR, 191-LEU, etc, as shown in Fig 9.

**Fig 9:**
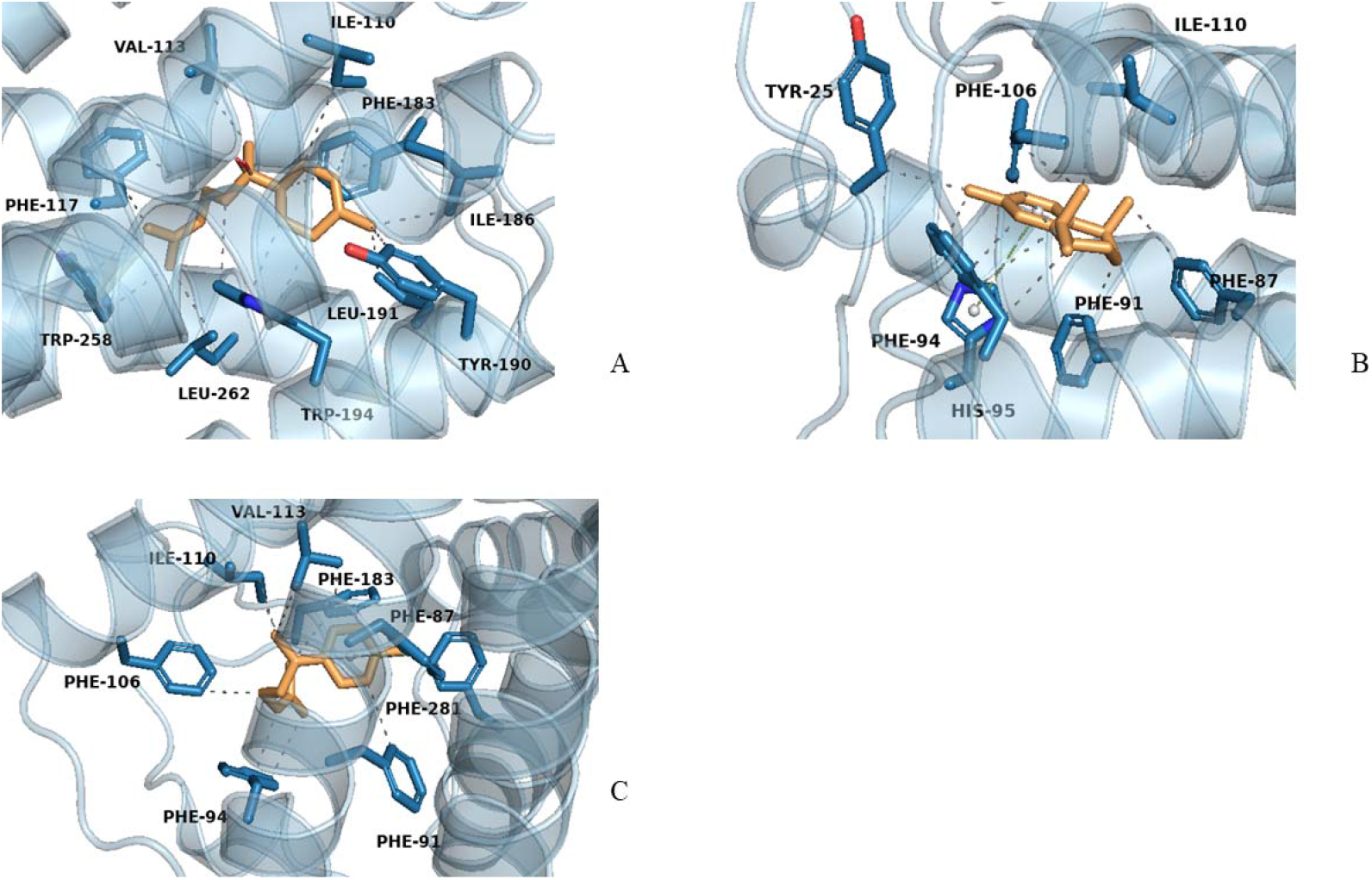
Protein ligand complexes formed by Bisabolol (A), Calamene (B) and α-Curcumene (C) with CNRII (PDB ID: 5ZTY)

When the molecular surface of the proteins was evaluated, it was seen that in order to reach the active site the small molecules have to ooze through a hole before binding, so if the molecule size in improper then the compound might not even reach the active site to bind. Out of all the docked complexes only the top performing ligand was simulated for further studies.

### Molecular Dynamic Simulation Studies

For reference purpose the apoprotein was also simulated along with the top-scoring terpene. CNR1 was compared with Gamma-Eudesmol and CNR2 was compared with Bisabolol in terms of RMSD, gyration and hydrogen-bond interaction. As we can see in Fig 10, CNR1 holds a steady RMSD median of 3.5□ throughout 100ns without any major fluctuations or unwanted spikes demonstrating complex instability. The CNR1-gamma-Eudesmol complex initially starts from a much lower fluctuation of almost 1□ and immediately jumps to almost 4□ and then it maintains a steady state RMSD throughout the simulation and finally merges with the apoprotein. All over the RMSD graph indicates high stability between the protein and the terpenoid, even from the radius of gyration graph it is evident that the complex formation stabilizes protein compactness (Supplementary Fig 3). To confirm the presence of Hydrogen bond interaction in Docking studies a simulated interaction was mapped out which shows that 1 Hydrogen bond was stable throughout the course of the simulation (Supplementary Fig 3).

**Fig 10:**
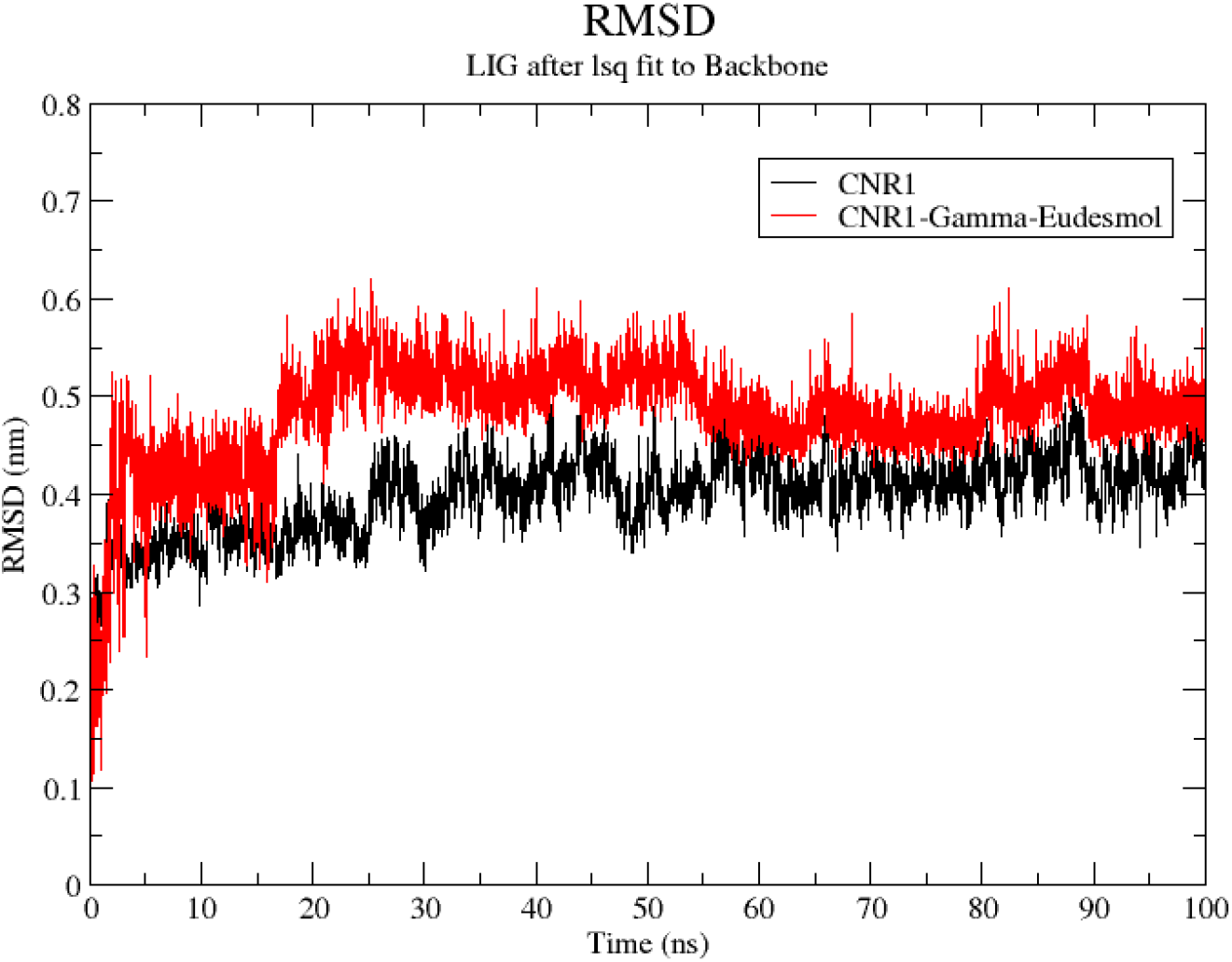
Molecular Dynamic simulation between the Apoprotein (Black) and the protein ligand complex (Red)

In case for CNRII due to unavailability of complete simulation results posed high RMSD values (almost 2□) due to greater fluctuations of unbound protein structure and missing residues from the protein structure (Fig 11). Interestingly when the ligand made a complex with protein the system stabilized and RMSD reduced subsequently even dropping to 0.5□ to a steady RMSD of 0.75□ as shown in Fig 11.

**Fig 11:**
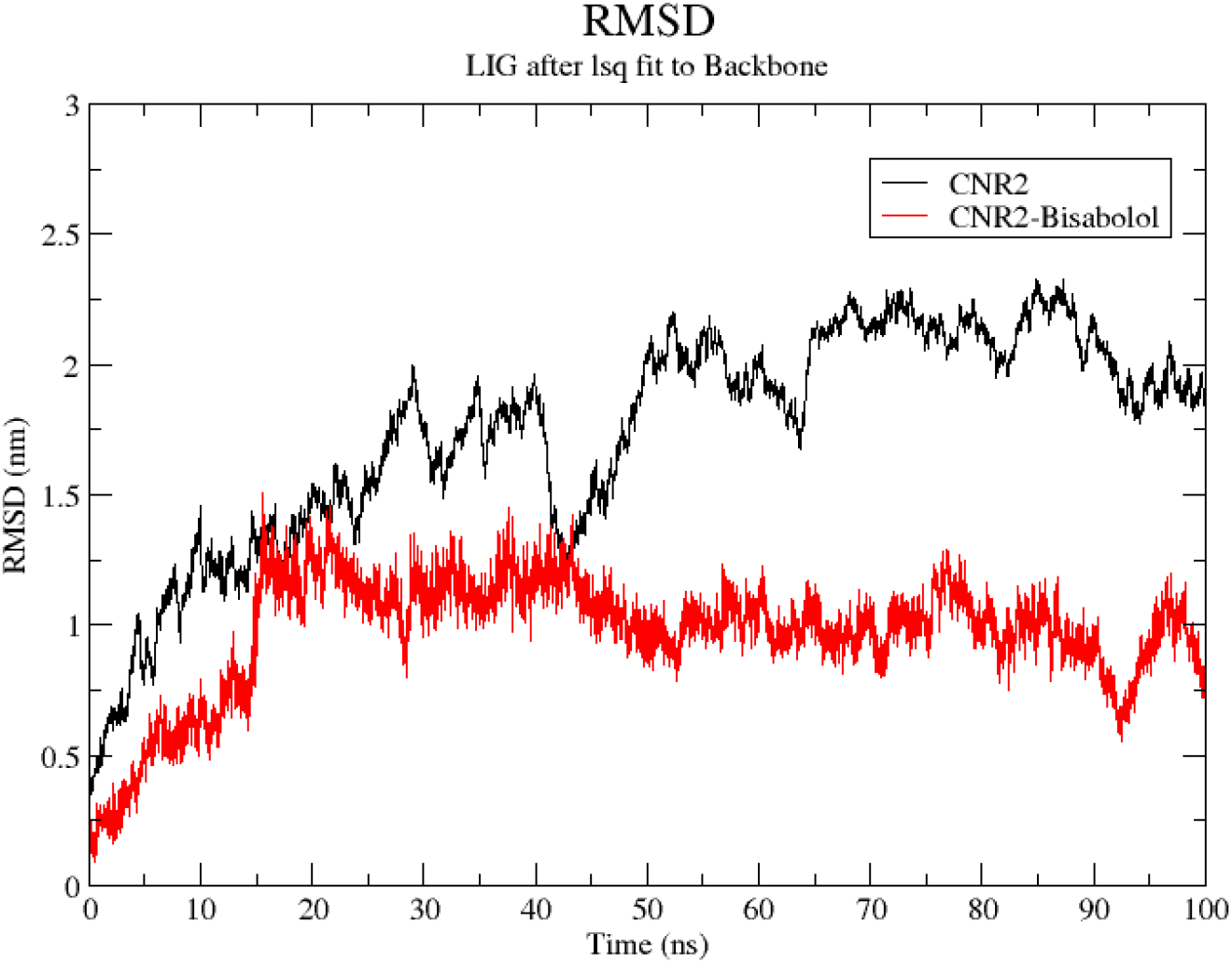
Molecular Dynamic simulation between the Apoprotein (Black) and the protein ligand complex (Red)

Even the radiation of gyration graph shows a steady compactness of the complex structure after 20ns whereas the radius of gyration of the apoprotein was quite higher and less stable than the complex (Supplementary Fig 4). No notable Hydrogen bond interactions were noted which substantiated the analysis from Molecular Docking. From the above representation of RMSD graphs, judging an explicit drug molecule might be a difficult task but both the systems have justified their roles in being a lead molecule for further studies up ahead.

### ADMET Analysis

Using SwissADME tools, the ADMET characteristics of two chosen compounds were thoroughly examined. The evaluation took into account factors such physiochemical characteristics, synthetic accessibility, drug-likeness (rule-based filtering), and bioavailability. The compounds’ ADMET profiles were visualised and compared using a heatmap and radar plot (Fig 12 and Fig 13).

**Fig 12:**
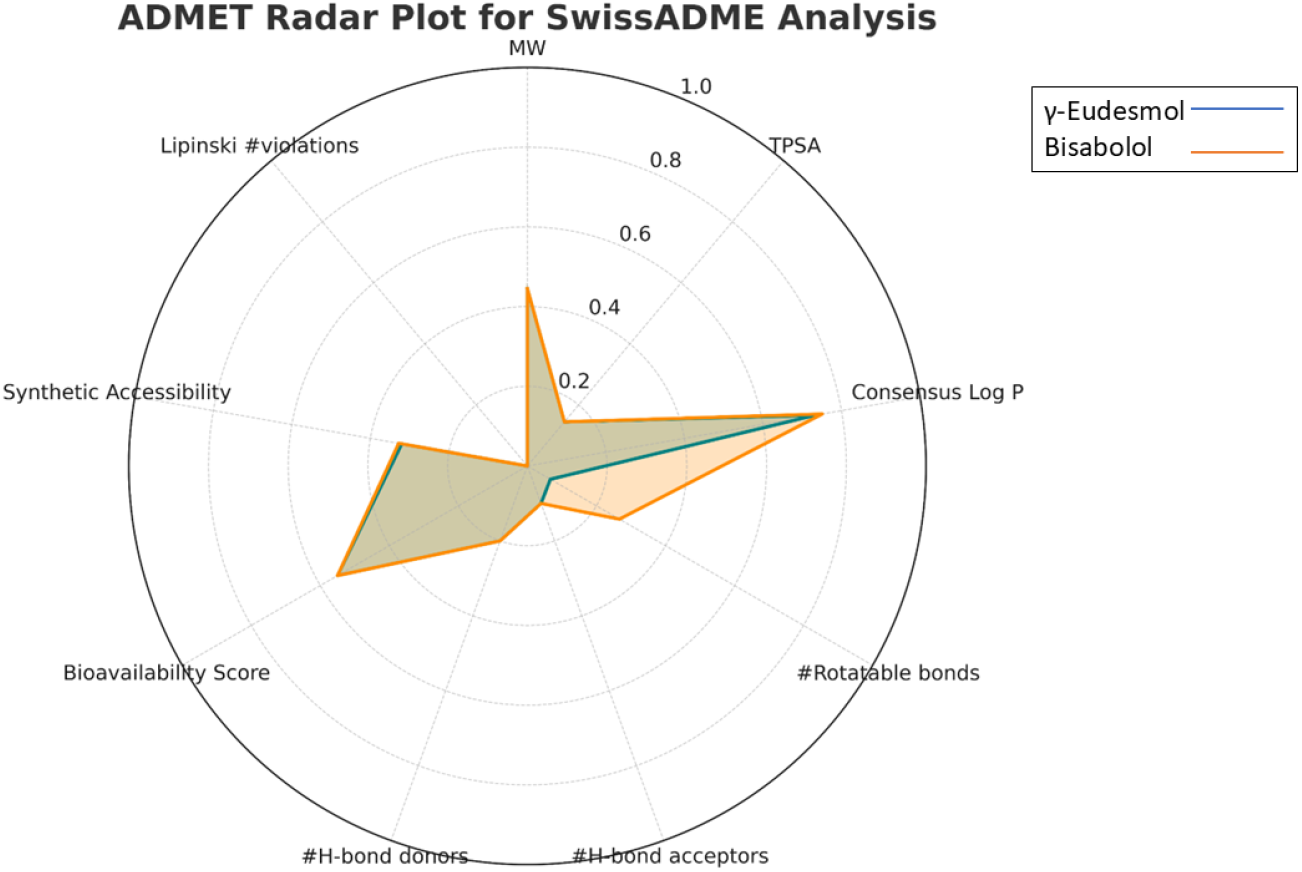
Radar Representation of γ-Eudesmol and Bisabolol

**Fig 13:**
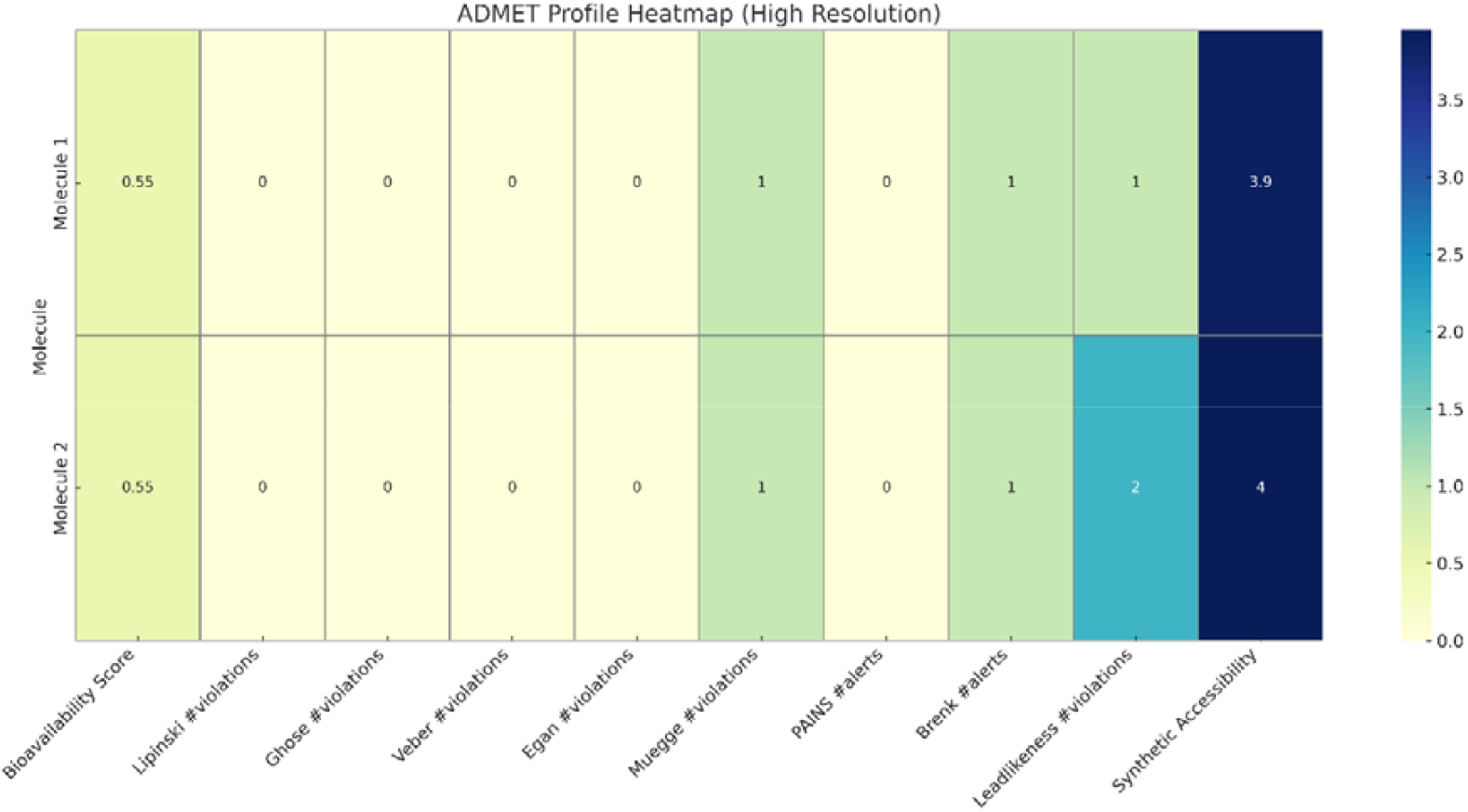
Heatmap Representation of γ-Eudesmol and Bisabolol

A visual comparison of normalised physiochemical factors that affect drug metabolism, membrane permeability, and oral bioavailability is shown in the ADMET radar map (Fig 12).

Notable findings include of:

- Comparable lipophilicity and membrane permeability are indicated by the two compounds’ similar consensus Log P values.
- Molecular weight (MW), topological polar surface area (TPSA), number of H-bond donors and acceptors, and number of rotatable bonds were all larger in Bisabolol. These characteristics point to increased solubility but maybe decreased passive permeability, which could have an impact on oral absorption.
- Although the difference is slight, Bisabolol’s bioavailability score is visibly higher, which is consistent with the heatmap data.
- γ-Eudesmol had a marginally better synthetic accessibility zone, which supports Fig 13’s finding.

Two candidate compounds’ synthetic feasibility metrics and rule-based drug-likeness filters are compiled in the ADMET heatmap (Fig 13).

Both γ**-Eudesmol** and **Bisabolol** exhibited:

- Strong adherence to established drug-likeness standards is confirmed by the fact that there were no infractions of the Lipinski, Ghose, Veber, Egan, or Muegge regulations.
- A moderate oral bioavailability score of 0.55 is generally regarded as acceptable in the early stages of drug research.

However, there were a few slight variations between the two:

- One PAINS alarm was generated by both compounds, indicating possible assay interference that should be taken into account during subsequent biological screening.
- Bisabolol displayed two lead-likeness violations, indicating greater structural complexity, whereas γ-Eudesmol exhibited one Brenk alert and one lead-likeness violation.
- γ-Eudesmol scored 3.9 on the synthetic accessibility scale, compared to Bisabolol’s 4.0, suggesting that γ-Eudesmol is slightly simpler to synthesise.

Although both compounds are drug-like and synthesizable, γ-Eudesmol seems more advantageous overall, according to the heatmap, because it has slightly fewer alarms and better synthetic feasibility.

While the radar plot (Fig 12) emphasises physicochemical characteristics pertinent to pharmacokinetics, the ADMET heatmap (Fig 13) places more emphasis on rule-based drug-likeness and synthetic feasibility. The conclusion that γ-Eudesmol exhibits a more favourable ADMET profile and merits prioritisation for additional in vitro and in vivo validation is supported by these visual aids taken together.

## Conclusion

By focussing on the cannabinoid receptors CNR1 and CNR2, this work used an integrative computational pipeline to investigate the medicinal potential of terpenoids synthesised from Cannabis sativa. Pharmacophore mapping, structure-based docking experiments, and ligand-based QSAR models were used to screen 119 structurally different terpenoids. In order to identify two lead pharmacophoric profiles, QSAR modelling for CNR1 and CNR2 produced statistically viable models (r2 = 0.854 and 0.798, respectively). After the terpenoid library was screened using these, γ-Eudesmol and Bisabolol were chosen as the best candidates for CNR1 and CNR2, respectively. Strong binding affinities of these terpenoids with their respective receptors were established by molecular docking; Bisabolol formed substantial hydrophobic contacts with the binding pocket of CNR2, while γ-Eudesmol displayed strong hydrogen bonding and hydrophobic interactions inside the CNR1 active site. Favourable RMSD trends and compactness (radius of gyration) across 100 ns simulations demonstrated that molecular dynamics simulations further confirmed the durability of these interactions. Both candidates demonstrated acceptable bioavailability and compliance with main drug-likeness requirements, according to ADMET analysis. However, γ-Eudesmol (Molecule 1) showed fewer structural alarms and somewhat greater synthetic accessibility than Bisabolol (Molecule 2). These results suggest that Bisabolol is still a strong contender for CNR2-associated pathways, while γ-Eudesmol is a promising lead chemical for the development of CNR1-targeted analgesics.

In conclusion, this study provides strong computational evidence for the potential analgesic applications of terpenoids and highlights their importance as viable scaffolds for cannabinoid receptor regulation. To confirm their pharmacological significance and biological activity, more in vitro and in vivo tests are relevant.

## Acknowledgements

The author would like to thank Central University of South Bihar Bioinformatics Department and Maulana Azad National Institute of Technology Biological Science and Engineering Department for constant support and
technical resources.

