## Supplementary material for "Insilico Investigation of Terpenoid efficacy on Cannabinoid Receptors using QSAR models and fragment-based Pharmacophore modelling": Suplementary

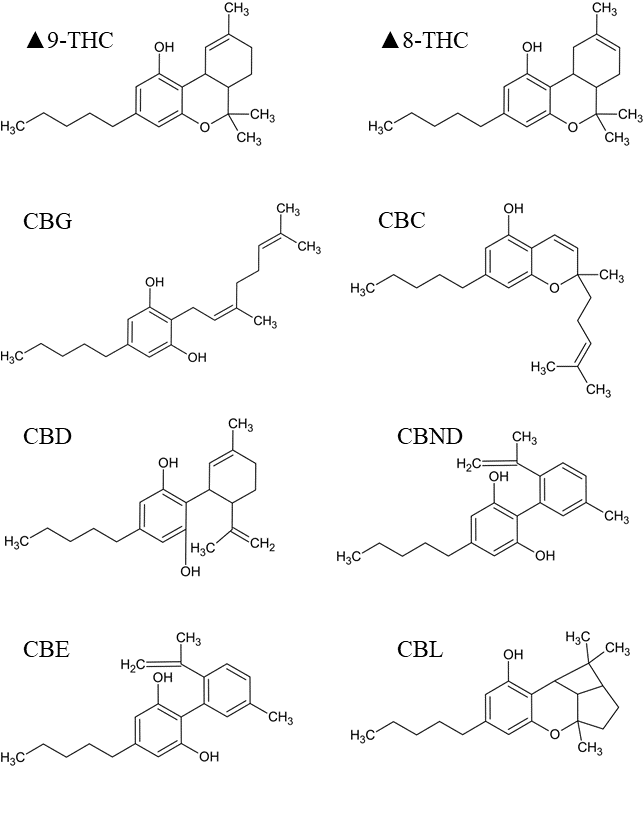

Supplementary Fig 1: Major Cannabinoids

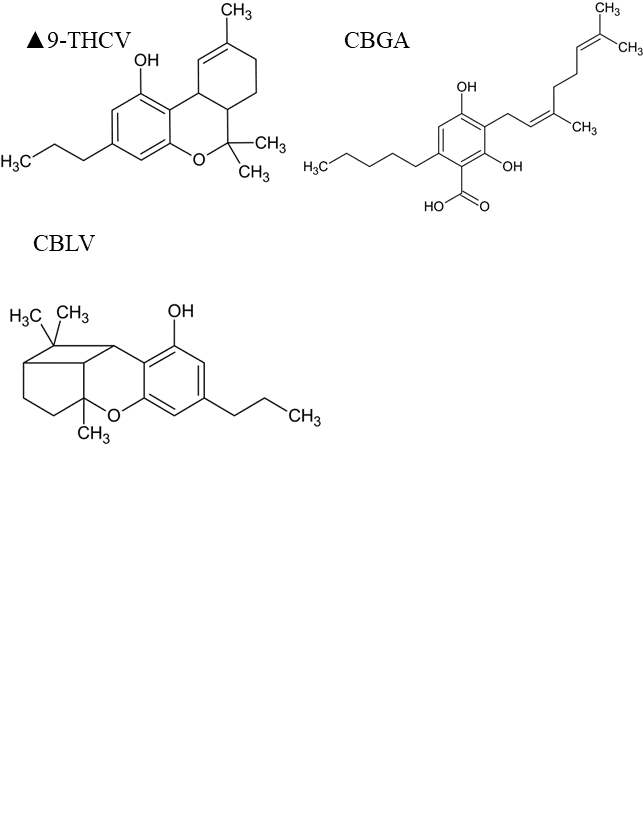

Supplementary Fig 2: Minor Cannabinoids obtained from *Cannabis sativa*

| **Terpenoid** | **PubChem CID** | **Terpenoid Class** |
| --- | --- | --- |
| p-cymene | 7463 | Mono Terpenes |
| dehydro-p-cymene | 62385 |  |
| myrcene | 31253 |  |
| limonene | 22311 |  |
| alpha-terpinene | 7462 |  |
| beta-phellandrene | 11142 |  |
| gamma-terpinene | 7461 |  |
| alpha-terpinolene | 11463 |  |
| alpha-pinene | 6654 |  |
| beta-pinene | 14896 |  |
| camphene | 6616 |  |
| linalool | 6549 |  |
| alpha-terpineol | 17100 |  |
| terpinene-4-ol | 11230 |  |
| linalool oxide | 6432254 |  |
| sabinene hydrate | 62367 |  |
| cis-beta- ocimene | 5320250 |  |
| trans-beta- ocimene | 5281553 |  |
| alpha-phellandrene | 7460 |  |
| delta-3-carene | 26049 |  |
| delta-4-carene | 78249 |  |
| sabinene | 18818 |  |
| alpha-thujene | 17868 |  |
| m-mentha- 1, 8-(9)-dien-5-ol | 118119770 |  |
| 2-methyl-2-heptene-6-one | 9862 |  |
| fenchyl alcohol | 15406 |  |
| borneol | 64685 |  |
| nerol | 643820 |  |
| geraniol | 637566 |  |
| carvacrol | 10364 |  |
| 1,8-cineol | 2758 |  |
| 1,4-cineol | 10106 |  |
| camphor | 2537 |  |
| 3-phenyl-2-methyl-prop-1-ene | 18687 |  |
| citral B | 643779 |  |
| citronellol | 8842 |  |
| geranyl acetone | 1549778 |  |
| carvone | 7439 |  |
| pulegone | 442495 |  |
| dihydrocarvone | 24473 |  |
| beta-terpineol | 8748 |  |
| dihydrocarveyl acetate | 30248 |  |
| p-cymene-8-ol | 14529 |  |
| beta-cyclocitral | 9895 |  |
| safranal | 61041 |  |
| cis-linalool oxide | 6428573 |  |
| perillene | 68316 |  |
| sabinol | 94147 |  |
| thujyl alcohol | 10550 |  |
| piperitone oxide | 92998 |  |
| piperitenone oxide | 442497 |  |
| fenchone | 14525 |  |
| bornyl acetate | 6448 |  |
| camphene hydrate | 101680 |  |
| alpha-pinene oxide | 91508 |  |
| pinocarveol | 102667 |  |
| pinocarvone | 121719 |  |
| ipsdienol | 92301 |  |
| cis-carveol | 330573 |  |
| cis-sabinene hydrate | 11744854 |  |
| alpha-humulene | 5281520 | Sesquiterpenes |
| beta-caryophyllene | 5281515 |  |
| caryophyllene oxide | 1742210 |  |
| curcumene | 92139 |  |
| alpha-trans-bergamotene | 86608 |  |
| alpha-selinene | 10856614 |  |
| beta-farnesene | 5281517 |  |
| longifolene | 289151 |  |
| humulene epoxide I | 5352470 |  |
| humulene epoxide II | 10704181 |  |
| caryophyllene alcohol | 11746218 |  |
| β-bisabolene | 10104370 |  |
| allo-aromadendrene | 42608158 |  |
| calamenene | 6429077 |  |
| α-copaene | 12303902 |  |
| nerolidol | 5284507 |  |
| alpha-gurjunene | 15560276 |  |
| iso-caryophyllene | 5281522 |  |
| beta-selinene | 442393 |  |
| selina-3,7(11)-diene | 522296 |  |
| selina-4(14),7(11)- diene | 10655819 |  |
| alpha-bisabolol | 1549992 |  |
| alpha-cedrene | 6431015 |  |
| alpha-cubebene | 442359 |  |
| delta-cadinene | 441005 |  |
| epi-beta- santalene | 91746538 |  |
| farnesol | 445070 |  |
| gamma-cadinene | 92313 |  |
| gamma-elemene | 6432312 |  |
| gamma-eudesmol | 6432005 |  |
| guaiol | 227829 |  |
| ledol | 92812 |  |
| trans-trans-alpha-farnesene | 5281516 |  |
| (Z)-beta-farnesene | 5317319 |  |
| farnesyl acetone | 1711945 |  |
| alpha-cadinene | 12306048 |  |
| alpha-cis-bergamotene | 6429303 |  |
| alpha-eudesmol | 92762 |  |
| alpha-guaiene | 5317844 |  |
| alpha-longipinene | 520957 |  |
| alpha-ylangene | 442409 |  |
| beta-elemene | 6918391 |  |
| beta-eudesmol | 91457 |  |
| epi-alpha-bisabolol | 1201551 |  |
| gamma-cis-bisabolene | 3033866 |  |
| gamma-curcumene | 12304273 |  |
| gamma-muurolene | 12313020 |  |
| gamma-trans-bisabolene | 5352437 |  |
| viridiflorene | 10910653 |  |
| germacrene-B | 15559495 |  |
| clovandiol | 76319362 |  |
| Phytol | 5280435 | Diterpenes |
| neophytadiene | 10446 |  |
| friedelan-3-one | 91472 | Triterpenes |
| epifriedelanol | 119242 |  |
| vomifoliol | 5280462 | Miscellaneous Terpenes |
| dihydrovomifoliol | 14135402 |  |
| beta-ionone | 638014 |  |
| dihydroactinidiolide | 27209 |  |

Supplementary Table 1: Terpenoid list with PubChem CID

| **PubChem CID** | **Inhibitory Constant Ki (in nM)** |
| --- | --- |
| 154969576 | 982 |
| 117688996 | 294 |
| 9966641 | 345 |
| 166594237 | 359 |
| 166594239 | 3.1 |
| 89995201 | 99 |
| 89994985 | 3.4 |
| 89994983 | 0.5 |
| 121283196 | 14.3 |
| 89994947 | 0.9 |
| 76310146 | 0.7 |
| 73387573 | 29.1 |
| 89994979 | 2.9 |
| 89994956 | 447 |
| 89994976 | 0.8 |
| 89994964 | 0.5 |
| 89994977 | 83.1 |
| 121283240 | 2.2 |
| 76317374 | 40.3 |
| 89994949 | 0.3 |
| 89994961 | 6.3 |
| 89994978 | 2.7 |
| 76317376 | 2.4 |
| 76310144 | 0.9 |
| 89994948 | 3.1 |
| 89994946 | 0.1 |
| 73330476 | 0.6 |
| 156027027 | 1.2 |
| 154406352 | 0.7 |
| 73387575 | 0.6 |
| 73387574 | 1.2 |
| 89994990 | 1 |
| 89995191 | 0.7 |
| 89994944 | 0.9 |
| 89994959 | 600 |
| 156027028 | 15.9 |
| 89994987 | 33.8 |
| 156027029 | 15.2 |
| 89994960 | 0.5 |
| 139295899 | 25.3 |
| 156027030 | 1 |
| 156027031 | 1 |
| 121289411 | 1 |
| 156027032 | 1 |
| 121283217 | 1000 |
| 89994969 | 2.2 |
| 121283233 | 2.4 |
| 73387577 | 3664 |
| 73387578 | 1000 |
| 73387579 | 941 |
| 89995208 | 1000 |
| 73387581 | 1000 |
| 73387583 | 1000 |
| 73387582 | 191 |
| 25030040 | 27 |
| 24896430 | 93.66 |
| 25030042 | 503 |
| 25030038 | 983.3 |
| 25029549 | 41.96 |
| 25029550 | 46.71 |
| 25029551 | 199 |
| 25029552 | 3.7 |
| 25029553 | 2.33 |
| 66941822 | 17.18 |

Supplementary Table 2: Cannabinoids with Biological Activity against Cannabinoid Receptor Type I

| **PubChem CID** | **Inhibitory Constant Ki (in nM)** |
| --- | --- |
| 25256778 | 1.66 |
| 25030040 | 2.94 |
| 25030041 | 4.77 |
| 24896430 | 1.07 |
| 46880678 | 0.27 |
| 25030042 | 2.32 |
| 25030037 | 454 |
| 25030038 | 159.09 |
| 25030039 | 224.98 |
| 25030043 | 1.75 |
| 25029549 | 4.7 |
| 25029550 | 2.3 |
| 25029551 | 1.4 |
| 25029552 | 81.95 |
| 25029553 | 5.69 |
| 66941822 | 37.18 |
| 154969576 | 40.5 |
| 117688996 | 33.1 |
| 9966641 | 28 |
| 166594237 | 12.9 |
| 166594239 | 0.8 |
| 89995201 | 802 |
| 89994985 | 1.2 |
| 89994983 | 1.4 |
| 121283196 | 1.2 |
| 89994947 | 1.8 |
| 76310146 | 0.4 |
| 73387573 | 8.7 |
| 89994979 | 7.7 |
| 89994976 | 0.6 |
| 89994964 | 1 |
| 89994977 | 36.8 |
| 121283240 | 7.1 |
| 76317374 | 8.2 |
| 89994949 | 0.7 |
| 89994961 | 2.8 |
| 89994978 | 1.5 |
| 76317376 | 1.5 |
| 76310144 | 1.4 |
| 89994948 | 4.5 |
| 89994946 | 0.2 |
| 73330476 | 0.8 |
| 156027027 | 0.8 |
| 154406352 | 0.8 |
| 73387575 | 0.7 |
| 73387574 | 0.6 |
| 89994990 | 0.8 |
| 89995191 | 1 |
| 89994944 | 2.7 |
| 89994959 | 1000 |
| 156027028 | 55.5 |
| 89994987 | 39.9 |
| 156027029 | 8.7 |
| 89994960 | 0.7 |
| 139295899 | 8.7 |
| 156027030 | 1 |
| 156027031 | 1 |
| 121289411 | 1 |
| 156027032 | 1 |
| 121283217 | 1000 |
| 89994969 | 6.2 |
| 121283233 | 6.5 |
| 73387577 | 94.8 |
| 73387578 | 1000 |
| 73387579 | 48.8 |
| 89995208 | 375 |
| 73387581 | 1000 |
| 73387583 | 700 |
| 73387582 | 20.9 |

Supplementary Table 3: Cannabinoids with Biological Activity against Cannabinoid Receptor Type II

| **PubChem CID** | **Ki (in nanomolar)** | **Experimental Log transformed Ki** | **Predicted Log Transformed Ki** |
| --- | --- | --- | --- |
| 25,029,550 | 46.71 | 7.33059 | 7.59458 |
| 89,994,949 | 0.3 | 9.52288 | 8.97563 |
| 76,317,374 | 40.3 | 7.39469 | 7.77165 |
| 121,283,233 | 2.4 | 8.61979 | 9.04493 |
| 121,283,217 | 1,000 | 6 | 6.1275 |
| 89,994,944 | 1 | 9.04576 | 9.43544 |
| 156,027,029 | 15.2 | 7.81816 | 7.46087 |
| 154,969,576 | 982 | 6.00789 | 6.47001 |
| 25,030,042 | 503 | 6.29843 | 6.32459 |
| 89,994,978 | 2.7 | 8.56864 | 8.15194 |
| 89,994,987 | 33.8 | 7.47108 | 8.31868 |
| 89,994,946 | 0.1 | 10 | 8.82402 |
| 89,994,976 | 0.8 | 9.09691 | 9.17914 |
| 156,027,028 | 16 | 7.7986 | 7.7642 |
| 89,994,985 | 3.4 | 8.46852 | 8.95295 |
| 121,283,196 | 14.3 | 7.84466 | 7.22873 |
| 24,896,430 | 93.66 | 7.02845 | 7.65319 |
| 89,994,959 | 600 | 6.22185 | 6.23278 |
| 25,030,040 | 27 | 7.56864 | 6.64272 |
| 89,994,956 | 447 | 6.34969 | 6.77591 |
| 89,995,201 | 99 | 7.00436 | 6.88472 |
| 156,027,031 | 1 | 9 | 8.54741 |
| 89,994,979 | 2.9 | 8.5376 | 9.03837 |
| 76,317,376 | 2.4 | 8.61979 | 8.16835 |
| 89,994,961 | 6.3 | 8.20066 | 8.40539 |
| 76,310,146 | 0.7 | 9.1549 | 9.22004 |
| 25,030,038 | 983 | 6.00731 | 6.35796 |
| 25,029,551 | 199 | 6.70115 | 7.2909 |
| 73,387,577 | 3,664 | 5.43604 | 5.96892 |
| 9,966,641 | 345 | 6.46218 | 6.77674 |
| 117,688,996 | 294 | 6.53165 | 5.82424 |
| 66,941,822 | 17.18 | 7.76498 | 7.768 |
| 89,994,960 | 0.5 | 9.30103 | 9.33727 |
| 25,029,549 | 41.96 | 7.37716 | 7.15413 |
| 89,994,947 | 0.9 | 9.04576 | 8.67682 |
| 25,029,552 | 3.7 | 8.4318 | 8.07108 |
| 25,029,553 | 2.33 | 8.63264 | 8.13007 |
| 73,330,476 | 0.6 | 9.22185 | 9.42573 |
| 73,387,582 | 191 | 6.71897 | 6.15733 |
| 89,994,948 | 3.1 | 8.50864 | 8.82402 |
| 73,387,581 | 1,000 | 6 | 6.21021 |
| 89,995,208 | 1,000 | 6 | 6.4373 |
| 121,289,411 | 1 | 9 | 8.50926 |

Table 4: Predicted Activity values on the base of QSAR modelling of active compounds on CNR I

| **PubChem CID** | **Ki (in nanomolar)** | **Experimental Log transformed Ki** | **Predicted Log Transformed Ki** |
| --- | --- | --- | --- |
| 89,995,201 | 802 | 6.09583 | 5.81742 |
| 25,029,550 | 2.3 | 8.63827 | 7.98461 |
| 76,317,374 | 8.2 | 8.08619 | 8.52594 |
| 73,387,579 | 48.8 | 7.31158 | 6.54219 |
| 121,283,233 | 6.5 | 8.18709 | 8.89216 |
| 121,283,217 | 1,000 | 6 | 6.76319 |
| 73,387,582 | 20.9 | 7.67985 | 7.01467 |
| 156,027,029 | 8.7 | 8.06048 | 8.17923 |
| 154,969,576 | 40.5 | 7.39254 | 7.69827 |
| 25,030,039 | 224.98 | 6.64786 | 7.21417 |
| 89,994,961 | 2.8 | 8.55284 | 7.87223 |
| 89,994,977 | 36.8 | 7.43415 | 7.77381 |
| 73,387,583 | 700 | 6.1549 | 6.87488 |
| 89,994,959 | 1,000 | 6 | 6.71807 |
| 9,966,641 | 28 | 7.55284 | 8.39734 |
| 156,027,028 | 55.5 | 7.25571 | 6.93504 |
| 89,994,976 | 0.6 | 9.22185 | 8.95771 |
| 121,283,196 | 1.2 | 8.92082 | 8.28153 |
| 24,896,430 | 1.07 | 8.97062 | 8.99563 |
| 89,994,944 | 2.7 | 8.56864 | 8.6951 |
| 25,030,038 | 159.09 | 6.79836 | 6.80453 |
| 76,317,376 | 1.5 | 8.82391 | 8.61081 |
| 89,994,979 | 7.7 | 8.11351 | 8.4447 |
| 89,994,990 | 0.8 | 9.09691 | 8.98854 |
| 156,027,031 | 1 | 9 | 8.97694 |
| 89,994,964 | 1 | 9 | 8.18537 |
| 73,387,581 | 1,000 | 6 | 6.12831 |
| 89,994,947 | 1.8 | 8.74473 | 8.5766 |
| 73,387,578 | 1,000 | 6 | 5.79563 |
| 25,030,037 | 454 | 6.34294 | 6.75293 |
| 25,029,551 | 1.4 | 8.85387 | 9.01789 |
| 46,880,678 | 0.27 | 9.56864 | 9.1896 |
| 89,994,985 | 1.2 | 8.92082 | 8.89172 |
| 117,688,996 | 33.1 | 7.48017 | 7.14702 |
| 25,030,040 | 2.94 | 8.53165 | 7.97752 |
| 89,994,946 | 0.2 | 9.69897 | 8.74025 |
| 25,029,549 | 4.7 | 8.3279 | 8.2203 |
| 76,310,144 | 1.4 | 8.85387 | 8.61081 |
| 25,029,552 | 81.95 | 7.08645 | 7.7186 |
| 25,029,553 | 5.69 | 8.24489 | 8.72997 |
| 25,030,041 | 4.77 | 8.32148 | 8.22553 |
| 73,387,574 | 0.6 | 9.22185 | 9.38988 |
| 76,310,146 | 0.4 | 9.39794 | 8.82101 |
| 73,387,573 | 8.7 | 8.06048 | 9.12897 |
| 89,994,983 | 1.4 | 8.85387 | 8.91203 |
| 121,289,411 | 1 | 9 | 8.95663 |

Table 5: Predicted Activity values on the base of QSAR modelling of active compounds on CNRII

| **PubChem CID** | **Fit Value** | **Mapped Features** |
| --- | --- | --- |
| 227829 | 3.17449 | HB_DONOR3, HYDROPHOBIC4, HYDROPHOBIC5, HYDROPHOBIC7 |
| 643820 | 3.06687 | HB_DONOR3, HYDROPHOBIC4, HYDROPHOBIC5, HYDROPHOBIC7 |
| 76319362 | 2.90602 | HB_DONOR3, HYDROPHOBIC4, HYDROPHOBIC5, HYDROPHOBIC7 |
| 445070 | 2.86988 | HB_DONOR3, HYDROPHOBIC4, HYDROPHOBIC5, HYDROPHOBIC7 |
| 8842 | 2.85397 | HB_DONOR3, HYDROPHOBIC4, HYDROPHOBIC5, HYDROPHOBIC7 |
| 6432005 | 2.79203 | HB_DONOR3, HYDROPHOBIC4, HYDROPHOBIC5, HYDROPHOBIC7 |
| 92762 | 2.40535 | HB_DONOR3, HYDROPHOBIC4, HYDROPHOBIC5, HYDROPHOBIC7 |
| 91457 | 2.22885 | HB_DONOR3, HYDROPHOBIC4, HYDROPHOBIC5, HYDROPHOBIC7 |
| 11746218 | 2.20174 | HB_DONOR3, HYDROPHOBIC4, HYDROPHOBIC5, HYDROPHOBIC7 |
| 6432254 | 1.97169 | HB_DONOR3, HYDROPHOBIC4, HYDROPHOBIC5, HYDROPHOBIC7 |
| 92812 | 1.69178 | HB_DONOR3, HYDROPHOBIC4, HYDROPHOBIC5, HYDROPHOBIC7 |
| 330573 | 1.54015 | HB_DONOR3, HYDROPHOBIC4, HYDROPHOBIC5, HYDROPHOBIC7 |
| 6429077 | 1.46607 | HYDROPHOBIC4, HYDROPHOBIC5, HYDROPHOBIC7, RING_AROMATIC13 |
| 92139 | 1.45499 | HYDROPHOBIC4, HYDROPHOBIC5, HYDROPHOBIC7, RING_AROMATIC13 |
| 119242 | 1.22099 | HB_DONOR3, HYDROPHOBIC4, HYDROPHOBIC5, HYDROPHOBIC7 |
| 10364 | 1.06748 | HB_DONOR3, HYDROPHOBIC4, HYDROPHOBIC5, HYDROPHOBIC7 |
| 5284507 | 1.00056 | HB_DONOR3, HYDROPHOBIC4, HYDROPHOBIC5, HYDROPHOBIC7 |
| 64685 | 0.734067 | HB_DONOR3, HYDROPHOBIC4, HYDROPHOBIC5, HYDROPHOBIC7 |
| 6428573 | 0.724879 | HB_DONOR3, HYDROPHOBIC4, HYDROPHOBIC5, HYDROPHOBIC7 |

Table 6: CID 6694182 pharmacophore - terpenoid library screening results

| **PubChem CID** | **Fit Value** | **Mapped features** |
| --- | --- | --- |
| 5284507 | 2.78929 | HB_DONOR7, HYDROPHOBIC10, HYDROPHOBIC12, HYDROPHOBIC13 |
| 10364 | 2.43927 | HB_DONOR7, HYDROPHOBIC12, HYDROPHOBIC13, RING_AROMATIC15 |
| 10364 | 1.71092 | HB_DONOR7, HYDROPHOBIC10, HYDROPHOBIC12, RING_AROMATIC15 |
| 92139 | 2.22037 | HYDROPHOBIC10, HYDROPHOBIC12, HYDROPHOBIC13, RING_AROMATIC15 |
| 6429077 | 2.02065 | HYDROPHOBIC10, HYDROPHOBIC12, HYDROPHOBIC13, RING_AROMATIC15 |
| 1549992 | 1.61766 | HB_DONOR7, HYDROPHOBIC10, HYDROPHOBIC12, HYDROPHOBIC13 |
| 92301 | 1.60048 | HB_DONOR7, HYDROPHOBIC10, HYDROPHOBIC12, HYDROPHOBIC13 |
| 5280435 | 0.981281 | HB_DONOR7, HYDROPHOBIC10, HYDROPHOBIC12, HYDROPHOBIC13 |
| 445070 | 0.823282 | HB_DONOR7, HYDROPHOBIC10, HYDROPHOBIC12, HYDROPHOBIC13 |
| 1201551 | 0.644448 | HB_DONOR7, HYDROPHOBIC10, HYDROPHOBIC12, HYDROPHOBIC13 |
| 11746218 | 0.383997 | HB_DONOR7, HYDROPHOBIC10, HYDROPHOBIC12, HYDROPHOBIC13 |

Table 7: CID 89994947 pharmacophore - terpenoid library screening results

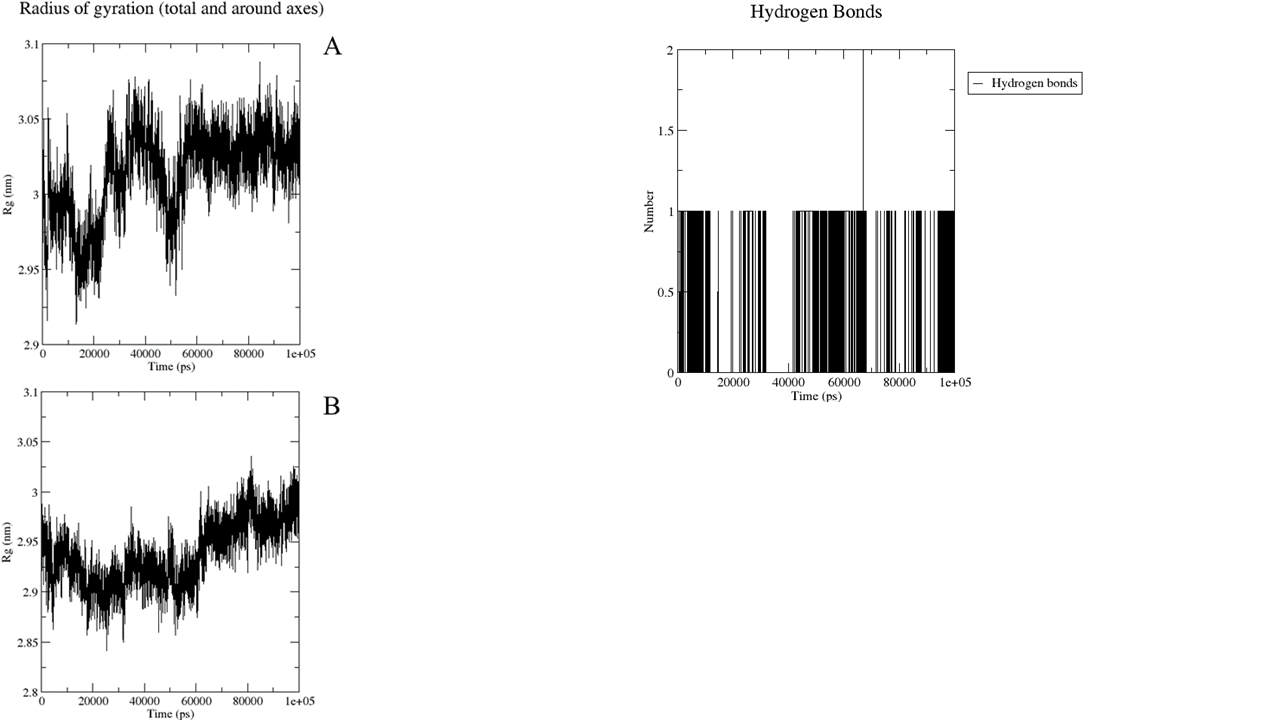

Supplementary Fig 3: Radius of Gyration of Apoprotein (A), Radiurs of Gyration of CNRI and γ- Eudesmol (Complex), Hydrogen Bond Interaction between CNRI and γ- Eudesmol (Complex)

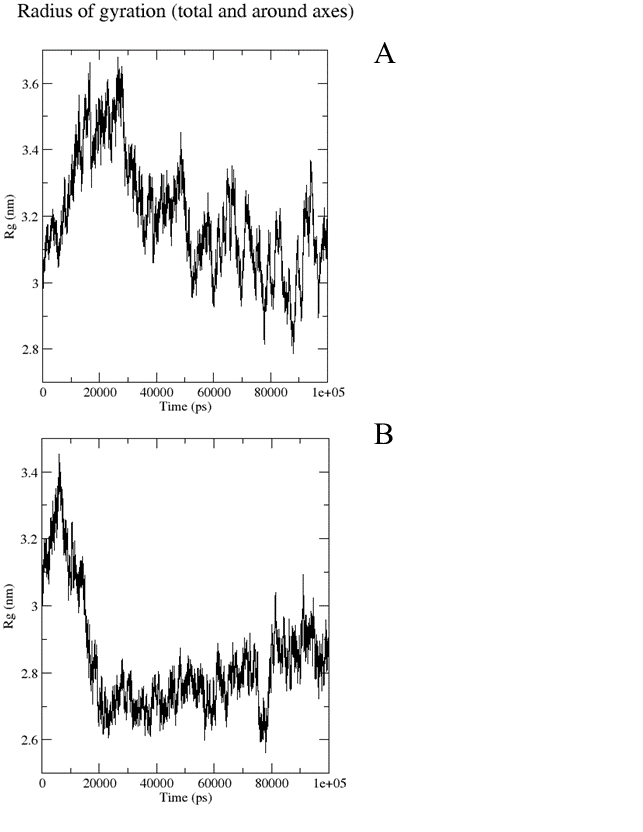

Supplementary Fig 4: Radius of Gyration of Apoprotein (A), Radiurs of Gyration of CNRII and Bisabolol (Complex)
